# Species-Specific ribosomal RNA-FISH identifies interspecies cellular-material exchange, active-cell population dynamics and cellular localization of translation machinery in clostridial cultures and co-cultures

**DOI:** 10.1101/2024.04.22.590569

**Authors:** John D. Hill, Eleftherios T. Papoutsakis

## Abstract

The development of synthetic microbial consortia in recent years has revealed that complex interspecies interactions, notably, the exchange of cytoplasmic material, exist even among organisms that originate from different ecological niches. Although morphogenetic characteristics, viable RNA and protein dyes and fluorescent reporter proteins have played an essential role in exploring such interactions, we hypothesized that <u>rRNA</u>-*fluorescence in <u>s</u>itu <u>h</u>ybridization* (rRNA-FISH) could be adapted and applied to further investigate interactions in synthetic or semisynthetic consortia. Despite its maturity, several challenges exist in using rRNA-FISH as a tool to quantitate individual species population dynamics and interspecies interactions using high-throughput instrumentation such as flow cytometry. In this work we resolve such challenges and apply rRNA-FISH to double and triple co-cultures of *Clostridium acetobutylicum, Clostridium ljungdahlii* and *Clostridium kluyverii.* In pursuing our goal to capture each organism’s population dynamics, we demonstrate the dynamic rRNA, and thus ribosome, exchange between the three species leading to formation of hybrid cells. We also characterize the localization patterns of the translation machinery in the three species, identifying distinct dynamic localization patterns among the three organisms. Our data also support the use of rRNA-FISH to assess the culture’s health and expansion potential, and here again our data find surprising differences among the three species examined. Taken together, our study argues for rRNA-FISH as a valuable and accessible tool for quantitative exploration of interspecies interactions, especially in organisms which cannot be genetically engineered or in consortia where selective pressures to maintain recombinant species cannot be used.

**IMPORTANCE:** Though dyes and fluorescent reporter proteins have played an essential role in identifying microbial species in cocultures, we hypothesized that <u>rRNA</u>-*fluorescence in <u>s</u>itu <u>h</u>ybridization* (rRNA-FISH) could be adapted and applied to probe, quantitatively, complex interactions between organisms in synthetic consortia. Despite its maturity, several challenges existed before rRNA-FISH could be used to study *clostridium* co-cultures of interest. First, species-specific probes for *Clostridium acetobutylicum* and *Clostridium ljungdahlii* had not been developed. Second, “state-of-the-art” labelling protocols were tedious and often resulted in sample loss. Third, it was unclear if FISH was compatible with existing fluorescent reporter proteins. We resolved key challenges and applied the technique to co-cultures of *C. acetobutylicum, C. ljungdahlii*, and *C. kluyveri.* We demonstrate that rRNA-FISH is capable of identifying rRNA/ribosome exchange between the three organisms and characterized rRNA localization patterns in each. In combination with flow cytometry, it can capture individual population dynamics in co-cultures.

## INTRODUCTION

Industrial microbiology has largely employed axenic culture strategies with selected wild-type or genetically engineered organisms or complex, naturally occurring microbial consortia, such as is the case in the dairy industry. For example, *Clostridium acetobutylicum* was grown historically in pure cultures to produce acetone and butanol from molasses before inexpensive petrochemicals made the process unprofitable in the mid 1900’s. The chemical industry is returning to fermentation as way to produce biofuels, chemicals, and bio-hydrogen more ecologically (1). In contrast to traditional approaches, synthetic microbial consortia, that is consortia consisting of two or more organisms that are not necessarily naturally co-existing, is a promising alternative (2, 3). Synthetic consortia offer some benefits over pure cultures such as division of labor, modularity, and the compartmentalization of incompatible metabolic reactions (2, 3). Using clostridial organisms, the co-culture approach for production of chemicals has attracted interest the last few years. Industrially interesting clostridia are categorized in groups based on their substrate and metabolic characteristics (4). Cellulolytic species can directly utilize lignocellulosics. Solventogens (an ill-defined term), such as *Clostridium acetobutylicum*, ferment carbohydrates into solvent molecules. Acetogens, such as *Clostridium ljungdahlii*, can fix CO_2_. Chain elongators, such as *Clostridium kluyveri*, convert short primary alcohols and carboxylic acids into longer-chain fatty acids. By combining species from different groups, the co-culture can be adapted to a variety of feedstocks, including syngas and lignocellulosic biomass, and produce desirable commodity chemicals and fuel molecules (1). In many cases clostridial organisms operate synergistically allowing for the production of novel metabolites with more efficient substrate conversion (5, 6).

Synthetic consortia provide a uniquely controlled environment to observe interspecies relations which may have been obscured by the inherent complexity of naturally occurring consortia. A variety of novel interspecies interactions have been revealed, including the formation of nanotube bridges, and cytoplasmic exchange mediated by cell fusion (3). Among those, cytoplasmic exchange is arguably the least expected. Benomar et al. first demonstrated that exchange of proteins between *Desulfovibrio vulgaris* and *C. acetobutylicum* can take place under nutritional-stress conditions and is mediated by cell-to-cell contact (7). Based on scanning electron micrographs, the cells remain physiologically distinct from one another during these events (7). Subsequent work emphasized the role of energetic coupling (i.e., contact mediated exchange of essential metabolites) between the two organisms and implicated the quorum sensing molecule autoinducer-II (AI-2) as the molecular basis of the interaction (8). Using anaerobic fluorescent proteins and protein dyes, our lab demonstrated heterologous-cell fusion driven cytoplasmic exchange between *C. acetobutylicum* and *C. ljungdahlii* (9). Unlike Benomar et al.’s system, *C. acetobutylicum* and *C. ljungdahlii* contact each other pole-to-pole, and that interaction can lead to heterologous, interspecies cell fusion and the formation of *hybrid cells*, cells that “contain uniformly distributed proteins and RNA from both organisms” (9). Most recently, it was shown that the cellular material exchange includes plasmid and chromosomal DNA exchange between *C. acetobutylicum* and *C. ljungdahlii* at frequencies comparable to transduction and conjugation in Gram-positive organisms (10). PacBio sequencing revealed that in some instances, *C. acetobutylicum* incorporated *C. ljungdahlii* plasmid and chromosomal DNA into its own chromosome (10). The mechanistic nature of the interaction and its implications on culture stability, especially on the long-term genetic stability, remain unexplored.

Does cytoplasmic exchange and interspecies cell fusion occur among other pairs of organisms? Of interest to our work are the interactions of *C. acetobutylicum* and *C. ljungdahlii* with *C. kluyveri*, the model organism which carries out chain elongation of linear carboxylic acids up to C_8_. Work characterizing the metabolic consequences of *C. acetobutylicum-C. kluyveri* and a *C. ljungdahlii*-*C. kluyveri* co-cultures has been published largely focusing on the metabolic potential of the co-cultures (6, 11). Our 2022 paper provides the first evidence for cytoplasmic material exchange between *C. acetobutylicum* and *C. kluyveri* (6). We sought to explore the interspecies relationships, especially cytoplasmic exchange, between these organisms. However, the transformation of *C. kluyveri* has yet to be reported in the literature. It cannot be made to express fluorescent proteins which has been essential for cytoplasmic tracking in the aforementioned studies. We hypothesized that *<u>r</u>ibosomal* <u>RNA</u>-*fluorescence in <u>s</u>itu <u>h</u>ybridization* (rRNA-FISH) could be used as an alternative marker for tracking rRNA exchange enabled by heterologous cell fusion. An rRNA-FISH probe is comprised of a short DNA sequence (16-40 bp) which bears homology to a target rRNA sequence. A fluorescent molecule, such as the Cy or Alexa Fluor family of fluorophores is attached to the 5’-end of the DNA probe. During hybridization, the probe forms a duplex with the target rRNA molecule, thus marking it fluorescently. rRNA-FISH has been applied to environmental microbiology since the 1990’s, where it is used to identify the presence of different taxonomic groups in naturally occurring consortia (12, 13). The DNA probes target sequences of the rRNA that are ideally unique to and ubiquitous among the targeted taxonomic groups. *Species-specific* rRNA-FISH is an embodiment of rRNA-FISH wherein the FISH probes bear homology to a species-specific region of the rRNA. Thus, individual species of bacteria can be identified among closely related organisms. Recently, it was applied to synthetic consortia to track the subpopulation dynamics of a co-culture containing *Clostridium carboxidivorans* (an acetogen) and *C. kluyveri.* Another important facet of rRNA-FISH in the context of synthetic consortia engineering is its unexplored potential to be used for assessing the overall culture health. Several studies have explored the relationship between cell viability and rRNA-FISH labelling, though in general, cell viability is difficult to determine for bacteria, especially in Gram-positive organisms (14). In *Escherichia coli*, rRNA-FISH was able to determine cell viability as well as other typical methods such as the BacLight Live/Dead and 5-Cyano-2,3-ditolyl tetrazolium chloride (CTC) assays when cells were exposed to UV light or heat, though all three methods overestimated the number of viable cells (15). rRNA-FISH has been assessed is overestimating the percentage of viable cells in a sample (15–17). In other words, rRNA-FISH tends to give false-positives, rather than false-negatives. Therefore, it cannot be used as a viability assay *per se*, though cell viability is a poorly defined parameter in general. However, unlabeled cells are almost certainly non-viable, as lacking a high concentration of rRNA (virtually all of which is bound in ribosomes, as we discuss below). We will examine this issue here using our data.

Here, we designed and validated novel *species-specific* rRNA-FISH probes for *C. acetobutylicum* and *C. ljungdahlii*. These probes target the 23S rRNA, the primary scaffold rRNA of the large ribosomal subunit. As the “state-of-the-art” hybridization procedure is complex and may lead to cell loss during processing, we aimed to simplify the protocol, removing in total twelve centrifugation and resuspension steps without any apparent loss of labelling efficacy. We validated that multiplexing with fluorescent protein reporters, such as HaloTag and protein dyes such as CellTracker dyes, is possible under the new protocol. In addition to tracking the individual species populations in clostridial co-cultures, we show that rRNA-FISH can assess a culture’s growth ability (or “health”). We demonstrate the potential of rRNA-FISH for identifying hybrid cells (i.e., cell containing rRNA from two different species) in a co-culture of *C. ljungdahlii* and *C. kluyveri* and a triple co-culture with *C. acetobutylicum,* thus enlarging the number of microbial pairs that interspecies exchange of cellular material takes place. Finally, we report the first rRNA/ribosome cellular localization studies in *C. acetobutylicum*, *C. ljungdahlii*, and *C. kluyveri*.

## RESULTS

### Validation of probe specificity and optimization of hybridization conditions

A species-specific rRNA-FISH probe set for *C. kluyveri,* ClosKluy, has previously been reported, but no suitable probes exist for *C. acetobutylicum* or *C. ljungdahlii* (18). Two sets of rRNA-FISH probes, ClosAcet and ClosLjun, were designed using the ARB (Latin, *<u>arb</u>or* meaning tree) software (19) and the SILVA (Latin, *<u>silva</u>* meaning forest) database (20) to target a species-specific region within the 23S rRNA of *C. acetobutylicum* and *C. ljungdahlii*, respectively. The specific probes sequences were chosen based on their predicted specificity in ARB, and because they had similar melting temperatures to ClosKluy (see Materials and Methods). We used an approach similar to Fuch et al. (21) to determine suitable hybridization conditions for ClosLjun, ClosAcet and ClosKluy. Formamide disrupts the hydrogen bonding which holds the DNA-RNA duplex together. Under optimal hybridization conditions, enough formamide is added to prevent imperfect duplex formation (off-target labelling), but overly stringent hybridization conditions prevent desirable bonding between probe and rRNA target. We tracked the median fluorescence of labelled *C. ljungdahlii, C. acetobutylicum*, and *C. kluyveri* at increasing formamide concentrations to determine the maximum stringency (i.e. maximum formamide concentration) that allowed strong labelling (Fig. 1A, 1B, and 1C). For instance, *C. acetobutylicum* was brightly labelled using ClosAcet at or below a formamide concentration of 20% (Fig. 1B). At higher formamide concentrations, the cells became less bright, suggesting that the increasing stringency was preventing ClosAcet from labelling *C. acetobutylicum* rRNA (Fig. 1B). *C. ljungdahlii* and *C. kluyveri* were separately hybridized with ClosAcet under the same conditions to demonstrate the absence of off-target labelling when using ClosAcet with these organisms (Fig. 1B). Analogous experiments were performed for the ClosLjun (Fig. 1A) and the ClosKluy probe sets (Fig. 1C). All three sets of probes demonstrated sufficient specificity and brightness at a formamide concentration of 20%.

**FIG 1.**
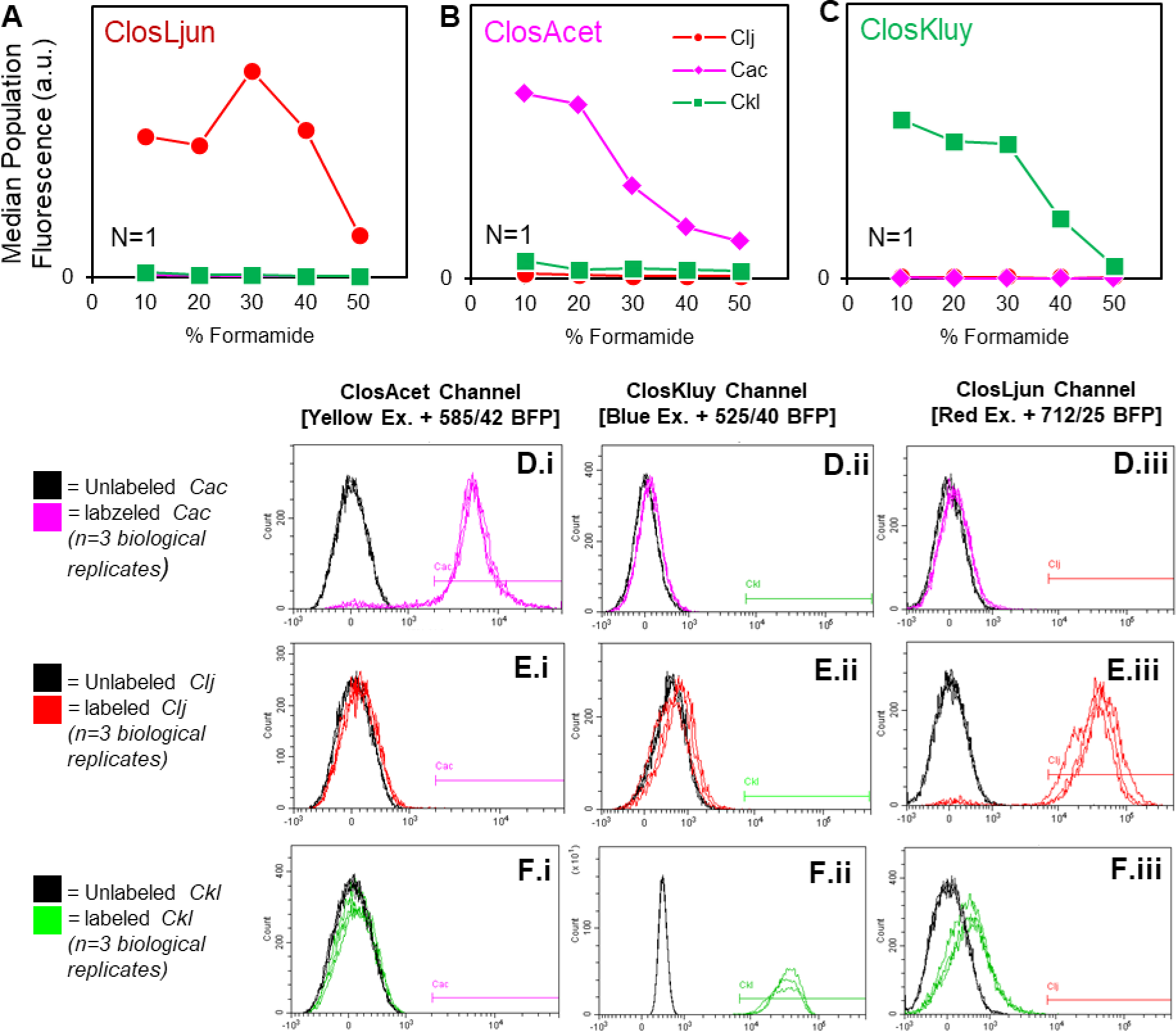
Optimization of formamide concentration and probe specificity. **(A-C)** *C. acetobutylicum* (Cac), *C. ljungdahlii* (Clj), and *C. kluyveri* (Ckl) were hybridized with each probe at formamide concentrations between 10% and 50% to determine ideal stringency. The median population fluorescence intensity (in arbitrary units, a.u.) was plotted for each sample on a linear axis indexed at 0. **(A)** ClosLjun selectively binds to *C. ljungdahlii’s* rRNA between 10-40% formamide, with optimal fluorescence at 30% formamide. **(B)** ClosAcet selectively binds to *C. acetobutylicum’s* rRNA between 10-20% formamide, with optimal fluorescence at 20% formamide. **(C)** ClosKluy selectively binds to *C. kluyveri’s* rRNA between 10-30% formamide, with optimal fluorescence at 10% formamide. **(D)** *C. acetobutylicum* was hybridized with ClosAcet, ClosKluy, and ClosLjun simultaneously at 20% formamide concentration and interrogated via flow-cytometry on the channels corresponding to each probe’s fluorescent marker. ‘labeled’ samples were compared to ‘unlabeled’ samples which underwent the same hybridization procedure but without any probes. **(E)** shows an analogous experiment to **(D)** performed in *C. ljungdahlii*. **(F)** shows an analogous experiment to **(D)** performed in *C. kluyveri*. For each species, only the corresponding probe induced a shift in the population’s fluorescence. The gating strategy established from this set of experiments was maintained throughout the work.

Further experimentation was necessary to ensure, first, the selectivity of the probes against off-target labelling, second, a high signal-to-noise ratio of the probes against autofluorescence, and third, the absence of bleed-over of the probe’s fluorescent signal into neighboring channels. Three biological replicates of each species were grown and sampled during exponential-phase growth, fixed, and incubated with ClosAcet, ClosLjun, and ClosKluy simultaneously under optimized hybridization conditions (20% formamide; discussed in the next section). These samples were compared to un-labelled negative controls to quantify autofluorescence. For all species, hybridization produced a strong shift in population level fluorescence when analyzed on the fluorescent channel corresponding to that species’ unique probe. For instance, labelled *C. acetobutylicum* (*Cac*) exhibited strong fluorescence on the ‘ClosAcet Channel’ when compared to the unlabeled control, indicating that autofluorescence is negligible (Fig. 1D.i). No such shift was observed for incompatible species/channel combinations, indicating the absence of bleed-over or off-target labelling. For instance, fluorescence of labelled *C. acetobutylicum* population overlapped with the negative control population when analyzed on the ‘ClosKluy Channel’ (Fig. 1D. ii) and the ‘ClosLjun Channel’ (Fig. 1D.iii) despite having been incubated with ClosKluy and ClosLjun. Further experimentation demonstrated the specificity of the probes against *C. ljungdahlii* (Fig. 1E.i, 1E.ii, and 1E.iii) and *C. kluyveri* (Fig. 1F.i, 1F.ii, and 1F.iii). Analogous confocal microscopy tests were performed on the same samples to validate the results obtained by flow cytometry (Fig. S1, S2, and S3). Fluorescence was only observed between corresponding species/channel combinations, but no fluorescent signal could be detected for unlabeled cells and incompatible species/channel combinations. Therefore, fluorescent signals appearing on microscopic images represent the presence of the targeted rRNA.

**FIG 2.**
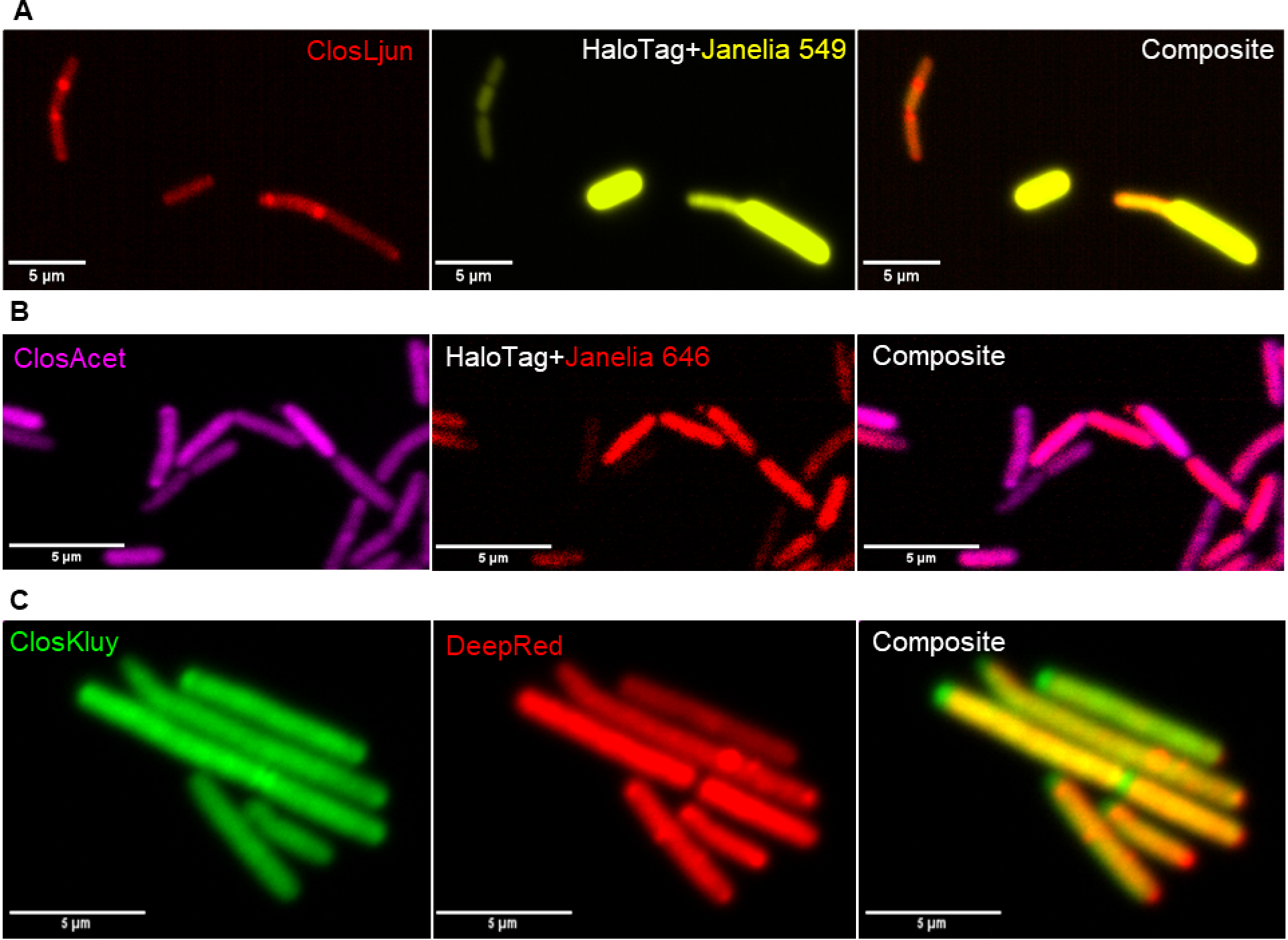
rRNA is compatible with common fluorescent protein labelling techniques. **(A)** C*. ljungdahlii*-p100ptaHalo (*Clj*-ptaHALO, top row) is labelled with ClosLjun and Janelia 549, a yellow emitting ligand for HaloTag. **(B)** *C. acetobutylicum*-p100ptaHalo (*Cac*-ptaHALO, bottom row) is labelled with ClosAcet (pseudo-colored magenta) and Janelia 646, a far-red ligand for HaloTag. **(C)** *C. kluyveri’*s proteins were labelled with CellTracker DeepRed since there have been no reports of successful exogenous gene expression in the organism. This dye was also found to be compatible with *in-solution* rRNA-FISH.

**FIG 3.**
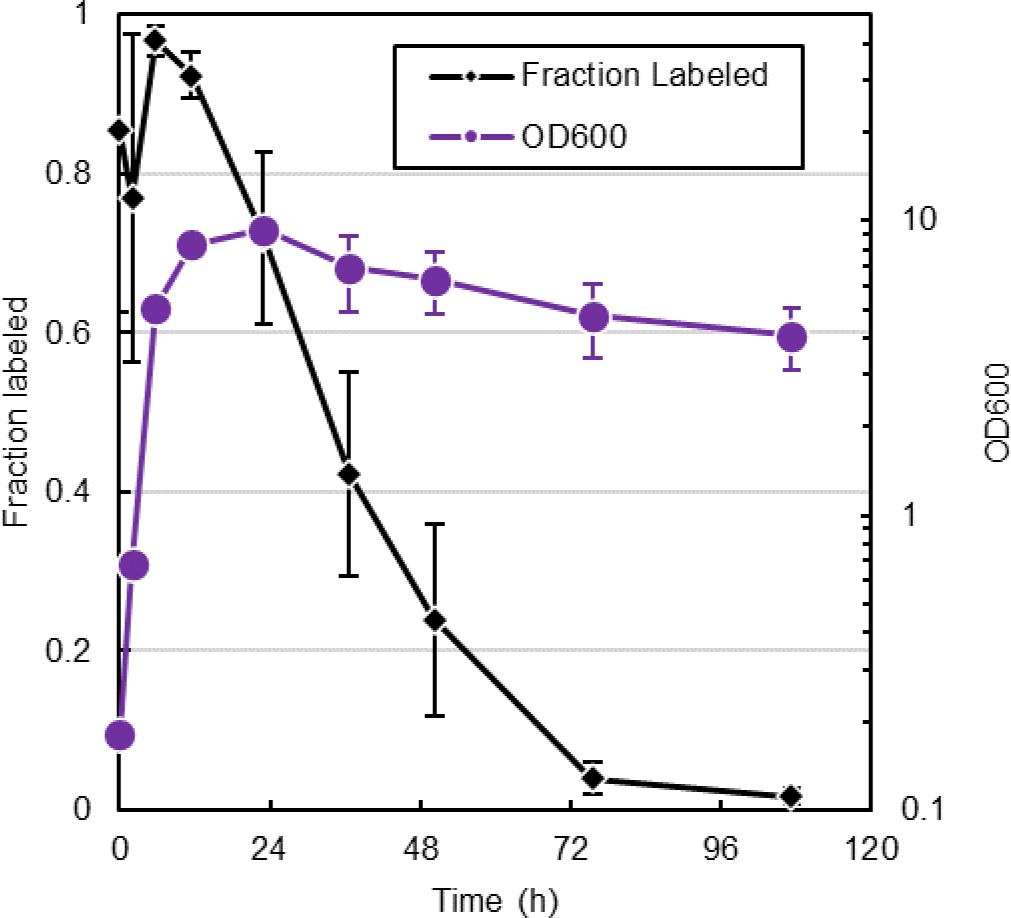
rRNA-FISH as an indicator of culture health/activity. OD_600_ and fraction of the population which was labelled by rRNA-FISH during batch cultivation of *C. acetobutylicum* in triplicate. Cells were labelled with ClosAcet. Cells with a fluorescent signal brighter than background were deemed “labelled.” Error bars represent a single standard deviation above and below the average.

Our rRNA-FISH probes target the 23S rRNA of the 50S large ribosomal subunit. Virtually all of the rRNA is incorporated into functional ribosomal subunits, as there is no significant pool of free rRNA (22). Only about 2-5% of rRNA is found in ‘ribosomal precursors,’ which are the immature ribonucleoprotein complexes which have yet to become a functional ribosome (23). Also, about 80% of ribosomes are actively translating (24). So, in essence, rRNA-FISH measures only bound rRNA, largely mature functional ribosomes, and thus rRNA-FISH fluorescence marks areas of active translation. While this notion has been challenged (25) for mixed environmental microbial populations due to the variable physiological characteristics of the diverse microbial populations in environmental samples, in pure cultures and defined synthetic consortia, published data (e.g., (26)) and our data below support this assertion.

### Optimizing rRNA-FISH for high-throughput flow-cytometric characterization

rRNA-FISH can be performed on slides or *in-solution*. *In-solution* rRNA-FISH is attractive because the sample can be analyzed using flow cytometry. When implemented this way, *species-specific* rRNA-FISH can be used to track subpopulation dynamics in co-culture (18). The major drawback to *in-solution r*RNA-FISH is that each treatment step requires centrifugation and resuspension which is time consuming and risks the loss of sample from incomplete centrifugation. Additionally, repeated centrifugation and incomplete resuspension could cause cell aggregation in the sample. The state-of-the-art approach for *in-solution* rRNA-FISH in *clostridium spp.* includes a fixation step (2-4 centrifugation/resuspension (C/R) steps), followed by dehydration (4 C/R steps), lysozyme treatment (2 C/R steps), further dehydration (4 C/R steps), hybridization, and washing steps (4 C/R steps) (27). During preliminary experiments, we discovered that many of the ‘canonical’ prehybridization steps for Gram-positive organisms are either not necessary or even detrimental in some cases. Fixation with paraformaldehyde (PFA) did not improve fluorescent signal compared to ethanol fixation, which is simpler and avoids the use of PFA, a known health hazard. Lysozyme treatment did not significantly improve signal in PFA fixed *C. acetobutylicum* and actually led to a complete loss of signal when fixed with ethanol (data not shown). In Bäumler et al.’s work, a dehydration step was included prior to hybridization as this was thought to better draw the probe in via osmotic forces (27, 28). We found that omitting the dehydration step prior to hybridization had no appreciable effect on any of the three organisms (Fig. S4A). Therefor all dehydration and the lysozyme permeabilization steps were omitted from our protocol, saving 12 C/R steps in total.

Hybridization conditions were screened to optimize the strength and speed of the probe binding in a manner similar to Wallner *et al.* using 20% formamide (29). We measured the time to maximum fluorescence for three probe concentrations: 0.1 μM (∼.61 ng/μL), 0.5 μM (3.1 ng/μL) and 1 μM (6.1 ng/μL). For each probe concentration and time point, cells from 3 biological replicates were sampled. The average of the three median population fluorescent intensity is plotted against time in Fig. S4B and Fig. S4C for *C. acetobutylicum* and *C. ljungdahlii,* respectively. Maximum fluorescence was reached between 3 and 5 hours, which is consistent with other reports of rRNA-FISH in firmicutes (30). We therefore conclude that the omission of prehybridization steps likely had a negligible effect on the rate of probe binding. 5 hours was sufficient to obtain a high fluorescent signal for both species with a probe concentration of 1 μM. Kinetic studies for *C. kluyveri* were omitted on the basis that previous work had reported success with 5 hour incubations (18).

### *In-solution* rRNA-FISH is compatible with anaerobic fluorescent reporter HaloTag and CellTracker Deep Red protein dye

Previously our lab has adapted the HaloTag fluorescent reporter protein system for use in *clostridium spp.* (31). HaloTag is a small protein (33 kDa) which is not fluorescent on its own but forms covalent bonds with a wide variety of fluorescent ligands (32). We verified that our newly developed rRNA-FISH labelling protocol is compatible with HaloTag fluorescence using HaloTag expressing strains: *C. acetobutylicum*-p100ptaHalo and *C. ljungdahlii*-p100ptaHalo (31). Cells from exponentially growing cultures were incubated with their respective “no-wash” HaloTag ligand prior to fixing and rRNA-FISH. For *C. acetobutylicum*-p100ptaHalo, Janelia 646, a far-red emitting ligand, was used since its fluorescent signal does not overlap with Cy3, the yellow emitting fluorophore of the ClosAcet probe set. For *C. ljungdahlii*-p100ptaHalo, Janelia 549, a yellow emitting ligand, was used since its fluorescent signal does not overlap with Cy5.5, the far-red emitting fluorophore of the ClosLjun probe set. Fig. 2A and Fig. 2B show the overlapping signal from both cells. HaloTag expression requires genetic modification which is not available in *C. kluyveri*. CellTracker Deep Red (Deep Red) has been used by our lab extensively to follow intercellular protein exchange (9) and forms covalent bonds with amine residues of proteins through a succinimidyl ester reactive group. To test if this too would survive *in-solution* rRNA-FISH, we labelled *C. kluyveri* with Deep Red prior to hybridization which resulted in strong double labelled *C. kluyveri* cells as seen in Fig. 3C. Together, we conclude that our *in-solution* rRNA-FISH protocol is compatible with other fluorescent labelling techniques which rely on covalent bonds.

### The *C. acetobutylicum* time-course of rRNA-FISH illustrates severe attenuation of translation in stationary phase due to commitment to sporulation and loss of growth potential

Previous work had determined that during batch cultivation of *C. acetobutylicum,* the number of colony-forming units (CFU) per volume decreases sharply once the culture reaches stationary phase (33). Since the OD_600_ and cell density remain constant during stationary phase, the decrease in CFU/mL has two potential sources: one is commitment to sporulation and the second is a shrinking viable cell population (34). As previously mentioned, rRNA-FISH labelling is correlated with active cell growth and translation, so we hypothesized that the labelled fraction of the population would decrease in later stationary phase as cells commit to sporulation and the viable fraction decreased. We performed three batch cultures of *C. acetobutylicum* and tracked OD_600_ and the fraction of the population which could be labelled by rRNA-FISH (Fig. 3). The labelled fraction sharply declined during at the onset of stationary phase, which corresponds to the increase of cells committing to sporulation and at the same time to a decrease in viable cell fraction which is typical of *C. acetobutylicum* batch cultivation. We would conclude that rRNA-FISH can be used to assess the growth (cell expansion) potential of a culture, but not strictly speaking, cell viability.

### *C. ljungdahlii* and *C. kluyveri* exchange rRNA and form rRNA hybrid cells

There have been several attempts to produce C_4_ to C_8_ alcohols and carboxylates from C_1_ gasses (e.g. CO and CO_2_) reported in the literature (35). Most were based on the co-cultivation of an acetogen (e.g. *C. ljungdahlii* (11), *C. autoethanogenum* (36), *Clostridium aceticum* (37), or *C. carboxidivorans* (27)) with *C. kluyveri*. In those systems, the acetogen consumes CO/CO_2_/H_2_ mixtures (syngas) to produce ethanol and acetate, which is further converted to butyric and caproic acid by *C. kluyveri*. Since the primary focus of such systems has been on metabolic production, interspecies interactions have been largely ignored. Diender et al. report that the presence of *C. kluyveri* redirects metabolic fluxes in *C. autoethanogenum*, resulting in increased ethanol production (36). The authors conclude that thermodynamic effects caused by continuous uptake of ethanol by *C. kluyveri*, contribute to the effect but do not go further.

Charubin et al. had hypothesized that heterologous cytoplasmic exchange was widespread in nature, having implicated syntrophic cross feeding as the primary impetus and recognized that syntrophic cross feeding is ubiquitous in naturally occurring microbiomes (9). Here, we hypothesized the *C. kluyveri* would be capable of forming hybrid cells with *C. ljungdahlii*, provided the culture conditions would create syntrophic interdependence. To do this, we modified the typical growth medium for *C. kluyveri* (TCGM-Ckl) which contains ethanol (343 mM) and acetate (166 mM) as the sole carbon sources. Acetate is an essential substrate of *C. kluyveri* when the only other carbon source is ethanol (38). Acetate is also the predominant product of energy metabolism in *C. ljungdahlii* when grown on fructose and gases, but *C. kluyveri* cannot use either of substrate for energy metabolism. For the co-culture, we modified TCGM-Ckl by removing acetate entirely from the medium, adding 11 mM fructose and pressurizing the headspace with 35-45 psi of H_2_/CO_2_. In so doing, *C. kluyveri* becomes dependent on acetate formation from *C. ljungdahlii*. We performed 3 biological replicates with a starting ratio of roughly 3:1 *C. ljungdahlii* to *C. kluyveri*. Samples were collected over 36 hours, labelled with ClosLjun and ClosKluy, and analyzed using flow-cytometry and microscopy. The growth of the culture (Fig. 4A), the relative sub-population fraction of each species (Fig. 4B), and the prevalence of “hybrid events” (Fig. 4C) were tracked. In this case, events with both ClosLjun and ClosKluy fluorescence constitute a ‘hybrid event,’ since the cell contains rRNA from both organisms. During the first 12 hours of growth, the OD_600_ roughly quadrupled, but the ratio of *C. ljungdahlii* to *C. kluyveri* remained relatively constant. At that point, the frequency of unidentifiable cells (‘unID’, Fig. 4B) increases sharply indicative of declining culture health. These correspond to cells that do not grow actively as they contain low concentrations of ribosomes/rRNA, thus displaying low translation activity. The ‘hybrid events’ have strong green fluorescence from ClosKluy and strong red fluorescence from ClosLjun. The frequency of ‘hybrid events’ reached its highest at 3.6 h, then decreased as culture health declined. The samples used for flow-cytometry were then examined by confocal microscopy. Select images from the first three time points are presented in Fig. 4D, 4E, 4F, and 4G. Cells with signal from both probes were found at roughly the frequency that flow cytometry predicted: approximately 1-2 cells on a random frame of about 100 cells.

**FIG 4.**
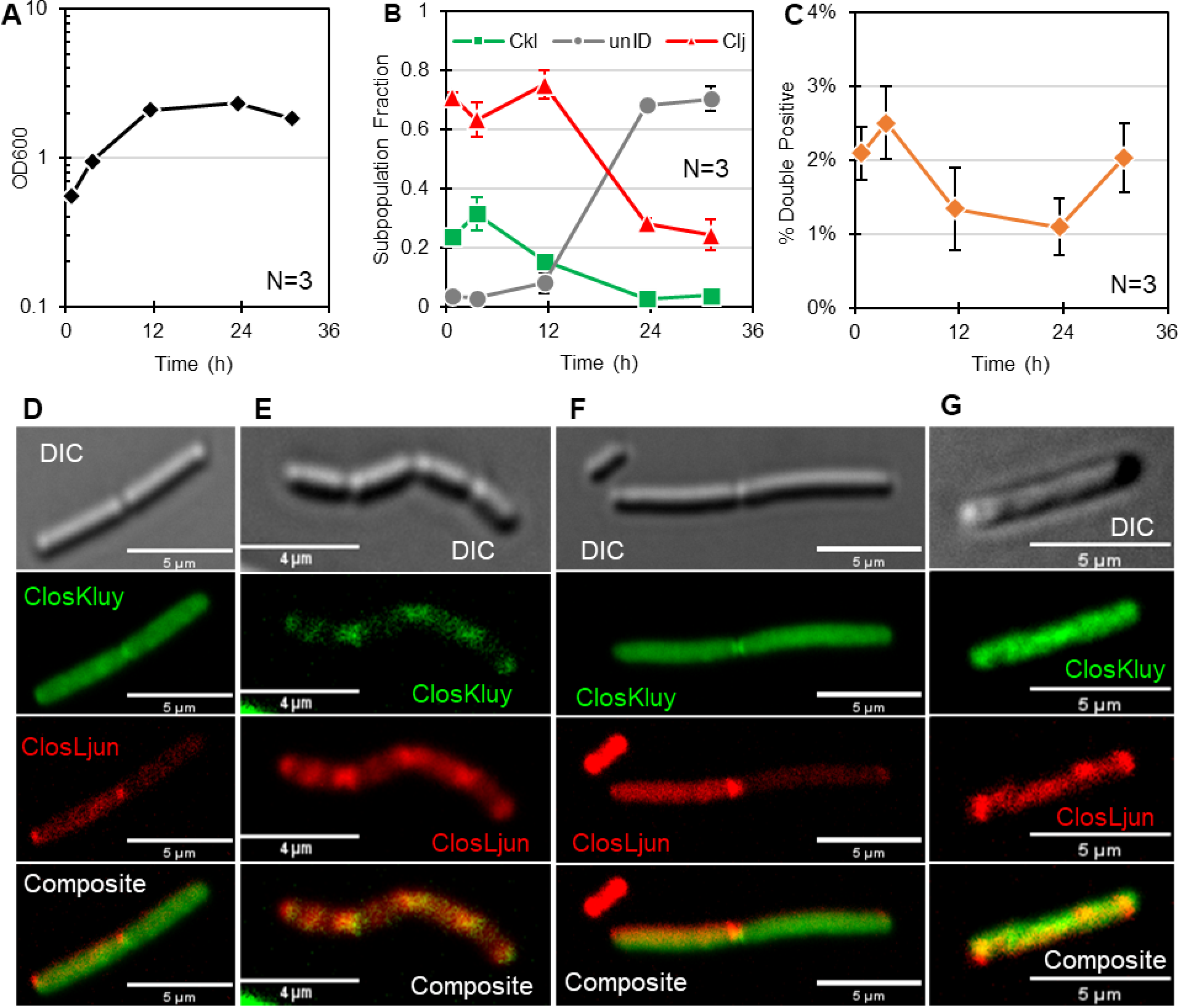
rRNA-hybrids in a co-culture of *C. ljungdahlii* (Clj) and *C. kluyveri* (Ckl). **(A)** The OD_600_ of three biological replicates. **(B)** The subpopulation fractions. **(C)** The hybrid cell frequency as a percentage of total. A hybrid cell from the 1-hour time point **(D)**, from the 4-hour time point **(E and F)** and from the 12-hour time point **(G)**.

### Revisiting the issue of protein and rRNA exchange between *C. acetobutylicum* and *C. ljungdahlii*

Charubin et al. had hypothesized that different macromolecules could be exchanged between *C. acetobutylicum* and *C. ljungdahlii* at different rates based on flow cytometric data that showed substantially faster (total) RNA exchange than protein exchange (9). Since the rRNA-FISH probes are compatible with fluorescent proteins, we were able to revisit this hypothesis by tracking the rate of protein and rRNA exchange simultaneously in a co-culture of *C. acetobutylicum* and *C. ljungdahlii*. A co-culture was performed between *C. ljungdahlii*-p100ptaHALO and *C. acetobutylicum* ATCC with an R-value of roughly 1.7, where the R-value is the starting ratio of *C. ljungdahlii* to *C. acetobutylicum* based on OD_600_ and preculture volume (5). Though *C. ljungdahlii-*p100ptaHALO precultures contained erythromycin, a translation inhibitor which targets the large ribosomal subunit, the cells were thoroughly washed prior to inoculation into the co-culture to prevent inhibition of *C. acetobutylicum* growth. Mono-culture controls were performed simultaneously (Fig. 5A). After the initial drop in pH caused by the accumulation of acids (Fig. 5B), the pH was maintained by addition of NaOH to prevent acid death (9). During the first 11 hours, samples were labelled with the HaloTag ligand OregonGreen, ClosAcet and ClosLjun, then analyzed via flow cytometry and microscopy (Fig. 5C, 5D, and 5E). OregonGreen was chosen as the fluorescent HaloTag ligand since its fluorescent signal does not overlap with ClosAcet (Cy3, yellow emitting, pseudo-colored magenta for clarity) and ClosLjun (Cy5.5, far-red emitting). *C. ljungdahlii* has two fluorescent labels, the HaloTag-OregonGreen complex and the ClosLjun probe (Fig. 5E). *C. acetobutylicum’s* cytoplasm is labelled only by the ClosAcet probe, (Fig. 5E). Population gating for HaloTag-OregonGreen signal was based on the mono-culture controls (Fig. S6). Population gating for ClosAcet and ClosLjun were based on the control experiments in Fig 1D and 1E and verified against mono-culture controls (Fig. S6). Either the coexistence of the ClosAcet and HaloTag-OregonGreen signals or the overlapping of the ClosAcet and ClosLjun signals would constitute a hybrid event. The largest fraction of hybrid events was found at 4 hours (Fig. 5C). At this time point, 2.65% of events exhibited ClosAcet/HaloTag-OregonGreen double fluorescence, while only 0.37% of events exhibited ClosAcet/ClosLjun double fluorescence. Other authors have pointed out that random cell aggregation may skew flow-cytometric results (27). Our data, however, demonstrate that hybrid events attributed to cell fusion are not artifacts of cell aggregation during culturing or labelling. If the ‘hybrid events’ were predominantly due to cell aggregates containing both species, we would expect those events to emit signals from ClosAcet, ClosLjun, and HaloTag-OregonGreen since the vast majority of cells attributed to the *C. ljungdahlii* population emit signals from both ClosLjun and HaloTag-OregonGreen (Fig. S7). The majority of hybrid events emit signal only from ClosAcet and HaloTag-OregonGreen and are, therefore, not aggregates. Moreover, aggregates are likely to form during repeated centrifugation and incomplete resuspension, but our simplified protocol obviates that risk. It is worth noting that microscopy images of hybrid cells, HaloTag fluorescence arose primarily from puncta representing probably several dozens to several hundreds of spatially associated HaloTag proteins (Fig. 5D). We hypothesize that these events primarily represent *C. acetobutylicum* cells which have acquired functional HaloTag proteins rather than *C. ljungdahlii* cells which have acquired *C. acetobutylicum* ribosomes based on the strong ClosAcet signal coming from these cells. Our findings are different from those of Charubin et al.’s findings that supported the hypothesis that RNA exchange occurs faster than protein exchange (9). This is likely due to differences in measurement techniques. Charubin et al.’s method labels, using a dye, total RNA and not only rRNA. As stated, fluorescence from rRNA-FISH represents assembled and largely actively translating ribosomes. Free mRNA and tRNA are much smaller molecules than ribosomes, and thus, compared to ribosomes, would be exchanged faster through cellular fusion events (9) due to enhanced molecular mobility.

**FIG 5.**
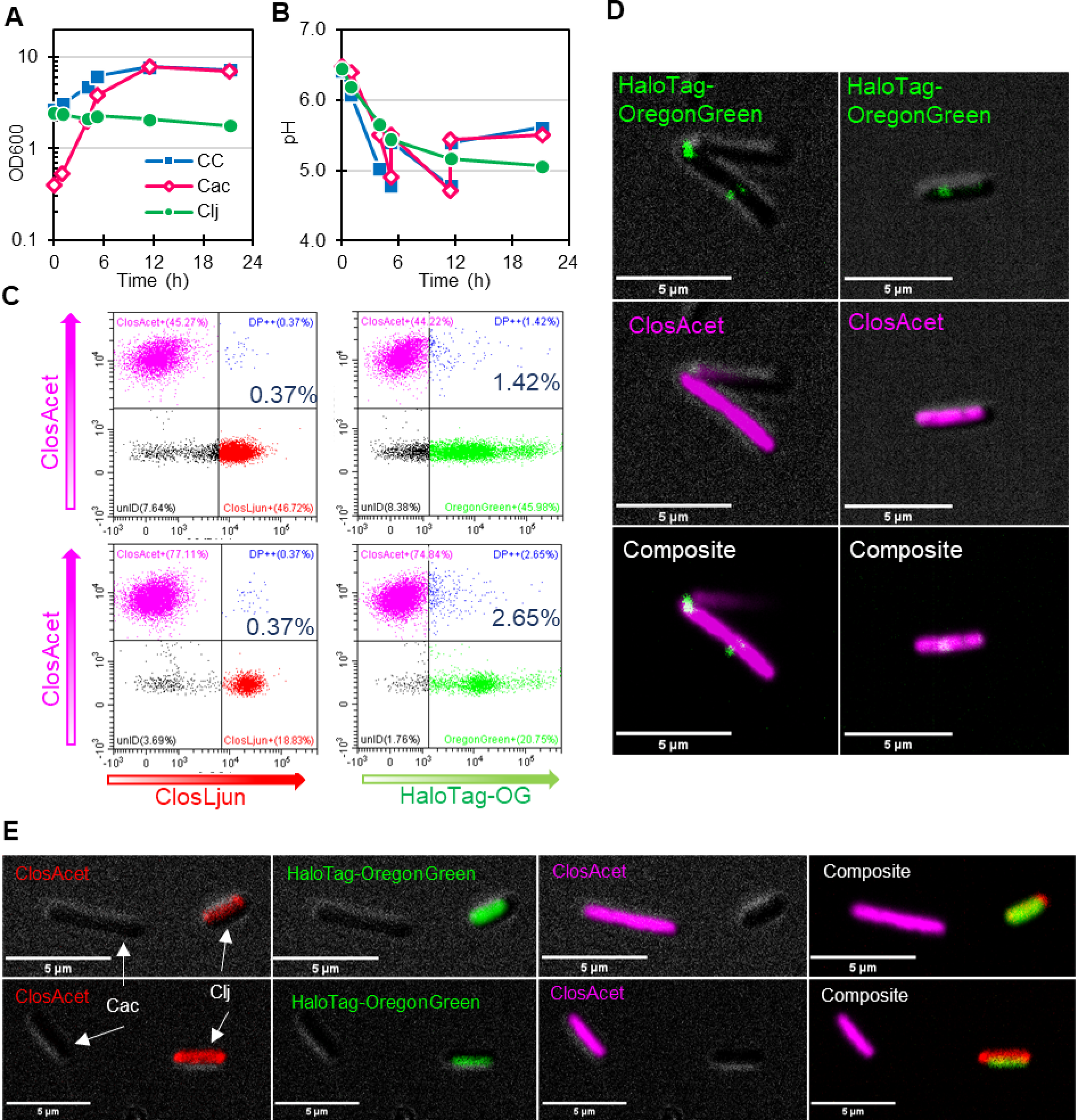
*C. ljungdahlii-*p100ptaHalo (Clj) to *C. acetobutylicum* (Cac) in co-culture. At 1, 4 and 11 hours, the culture was sampled and labelled with ClosAcet, ClosLjun, and HaloTag-OregonGreen (HaloTag-OG). **(A)** The OD_600_ and **(B)** pH of the co-culture and mono-culture controls. **(C)** Flow-cytometry dot plots measured the frequency of double positives with signals from ClosAcet and either HaloTag-OregonGreen (right column) or *C. ljungdahlii’s* ClosLjun labelled rRNA (left column). **(D)** Fluorescent microscopy images of hybrid cells from the 4-hour timepoint containing rRNA from *C. acetobutylicum* (magenta) and HaloTag-OregonGreen (green) from *C. ljungdahlii*-p100ptaHALO. HaloTag-OregonGreen fluorescence forms puncta. For comparison, **(E)** shows confocal microscopy images of the non-hybrid phenotype from the same time point. Dim DIC images are provided in the background to show cell location. *C. acetobutylicum* exhibits strong ClosAcet (magenta) fluorescence. *C. ljungdahlii-*p100ptaHalo exhibits both green and red signal from the HaloTag-OregonGreen complex and ClosLjun, respectively.

### rRNA FISH tracks the population dynamics of a triple coculture and identifies binary fusion events

To demonstrate the multiplexing ability of ClosAcet, ClosKluy, and ClosLjun, we applied our hybridization protocol to a synthetic consortium of *C. acetobutylicum, C. ljungdahlii*, and *C. kluyveri* and mono-culture controls. The optical density of the cultures is shown in Fig. 6A, and the sub-population dynamics for the triple co-culture are shown in Fig. 6B. In this case, the preculture and culturing conditions were such that *C. acetobutylicum* were suboptimal, and this could be determined by rRNA-FISH. There is agreement between the growth behavior of the mono-cultures and those in the triple culture. Both *C. kluyveri* and *C. ljungdahlii* grew substantially in the mono-cultures, but *C. acetobutylicum* did not. This pattern is reflected in the subpopulation dynamics, insomuch as *C. kluyveri* and *C. ljungdahlii* become the predominant species and *C. acetobutylicum* becomes the minor population. The rRNA-FISH data also suggests large numbers of non-actively translating cells, indicated by the large unidentifiable population (i.e. “unID” in Fig. 6B). In healthy, exponentially growing cultures, the fraction of unlabeled cells rarely rises above 10% (discussed in next section), but the fraction of unlabeled cells in this culture was roughly 40% during growth.

**FIG 6.**
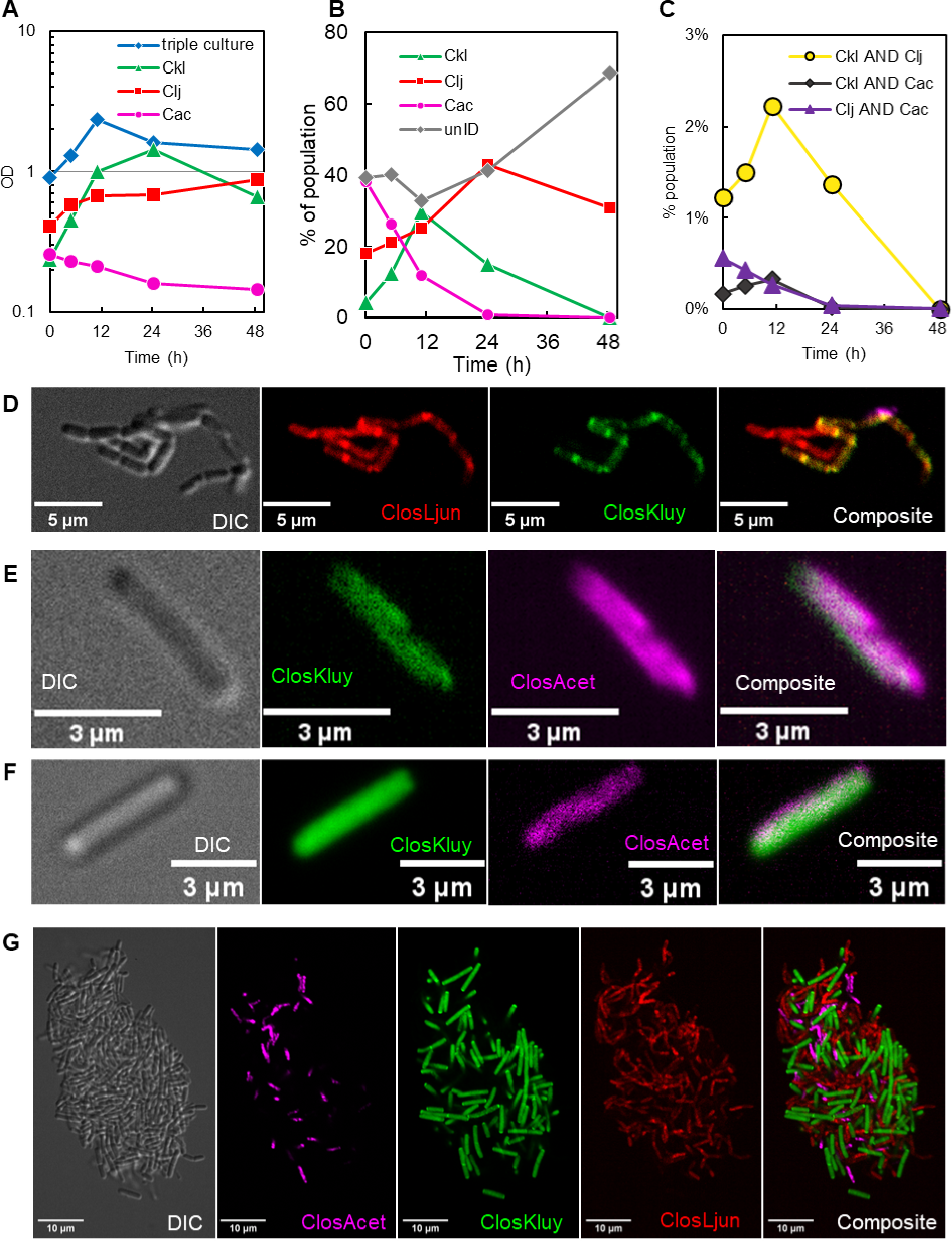
rRNA-FISH applied to a triple culture of *C. acetobutylicum*, *C. kluyveri*, and *C. ljungdahlii* and mono-culture controls. **(A)** The optical density of the triple culture and mono-culture controls. **(B)** The subpopulation dynamics of the three organisms and unidentifiable cells (“unID”) which are deemed inactive since they lack sufficient translation machinery to be clearly labelled by rRNA-FISH. **(C)** The frequency of the hybrid populations as measured by flow-cytometry. **(D)** A cluster of cells containing *C. ljungdahlii*-*C. kluyveri* hybrid cells. **(E and F)** *C. acetobutylicum*-*C. kluyveri* hybrids. **(G)** A biofilm fragment containing all three species isolated from the culture.

We also tracked the prevalence of the binary hybrid populations via flow-cytometry (Fig. 6C). *C. kluyveri*-*C. ljungdahlii* hybrids were more common than hybrids containing *C. acetobutylicum*. This may be in part due to the poor health of *C. acetobutylicum,* but the motivations and mechanism of heterologous cell fusion is not fully understood. Microscopy was performed to capture the hybrid events (Fig. 6D, 6E, and 6F). A cluster of *C. kluyveri*-*C. ljungdahlii* hybrids is presented in Fig. 6D. In this cluster, nearly all cells contain the ClosLjun probe, but one chain also contains the ClosKluy probe. This suggests that the green fluorescence is not merely a fluorescent artifact, but rather that it comes from ClosKluy-labelled *C. kluyveri* rRNA that co-exists with *C. ljungdahlii* rRNA. Since all cells in this cluster and neighboring cells experienced an identical hybridization environment and handling, we would expect to see identical labelling if *C. kluyveri* rRNA was not actually present. Moreover, the spatial pattern of ClosKluy and ClosLjun labelling is nearly identical suggesting that the translation machinery from both organisms is localized in the same cellular compartments. It is possible that rRNA from both organisms form actively translating ribosomes. The peculiarity of morphology prompted the following section’s investigation into the characteristic ribosomal localization patterns of each organism. Fig. 6E and Fig. 6F present *C. acetobutylicum*-*C. kluyveri* hybrids which were more difficult to find. These events are similar to those presented in previous work from our group (6). Finally, a small biofilm fragment was isolated from the labelled culture and imaged to demonstrate the potential of rRNA-FISH to deconvolute complex interspecies structures (Fig. 6G).

### Characterization of cellular rRNA localization in *C. kluyveri, C. acetobutylicum*, and *C. ljungdahlii*

Unlike well studied microorganisms such as *E. coli* and *B. subtilis*, subcellular organization has not been extensively studied in *C. kluyveri*, *C. acetobutylicum*, or *C. ljungdahlii*. In many prokaryotic species, the nucleoid, which is the area of the condensed chromosome and the transcription machinery, occupies the center of the cell and excludes the translation machinery. rRNA localization studies of model organisms has been an ongoing endeavor since about 2000 (39). An important characteristic is the ratio of the nucleoid size to the cytoplasm size which is known as the N/C ratio. Generally, the nucleoid size does not scale proportionally with cell size, in that the nucleoid of a larger cells occupies a smaller portion of the cytoplasm (i.e. larger cells have smaller N/C ratios) (40). In *E. coli*, the strength of nucleoid exclusion was inversely related to the N/C ratio (40). From this, the authors concluded that larger cells, such as firmicutes (which have typically low N/C ratios) would tend to exhibit stronger exclusion of translation machinery (40). Indeed, *B. subtilis*, the model organism for firmicutes, exhibits strong nucleoid exclusion effects (26, 41). Our rRNA-FISH probes target the 23S rRNA of the 50S large ribosomal subunit. Virtually all of the rRNA is incorporated into functional ribosomal subunits. Only about 2-5% of rRNA is found in ‘ribosomal precursors,’ which are the immature ribonucleoprotein complexes which have yet to become a functional ribosome (23), and as stated previously, there are no free rRNA in the cells (22). That the transcription and translation machineries are separated apparently challenges the well-established paradigm in microbial biology that translation initiation occurs on the nascently forming mRNA transcript. It is important to distinguish between a “bound” ribosomes which are a complex of a large and small subunit, mRNA, and the nascent polypeptide chain, and “unbound” ribosomes which are simply the individual subunits. About 80% of ribosomes are actively translating (24), and this is important in light of work which demonstrates that while the majority of translation occurs outside the nucleoid, unbound ribosomal subunits freely diffuse throughout the cell (42). One would conclude that compartmentalized rRNA-FISH fluorescence represents the compartmentalization of actively translating ribosomes.

We observed the most distinct ribosomal localization and striking division behavior in *C. ljungdahlii* (Fig. 7). *C. ljungdahlii* maintained exponential growth for about 20 hours after inoculation, during which fluorescence was high at the population level (Fig. 7A and 7B). At OD_600_ of 0.3, corresponding to t3 (Fig. 7A), most cells formed elongating ‘chains’ of replicating but not separating cells. DIC images clearly show the nascent cleavage furrow forming throughout the chain at regular intervals (Fig. 7C, Fig. S11). For each microscopic image, the full frame image is provided in the supplemental figures to support the argument that the cells provided in the main figures are representative/typical for the entire population. The plots to the right of the microscopy images present the fluorescent intensity along the major axis of the bacilliform-cell chain going from left to right. The fluorescent intensity is plotted in arbitrary units so as to compare the relative fluorescence (i.e. the prominence) of the signal within a cell, but not between cells. Microscopy of stationary phase *C. ljungdahlii* (Fig. S12), confirms the expected, canonical drop in fluorescence (Fig. 7B), which here, given the lack of *C. ljungdahlii* sporulation under these culture conditions, is likely caused by carbon starvation and a rapid decrease in the number of viable cells. One of the few fluorescent cells at the t5 stationary-phase point is shown in Fig. 7D, but the prominence of the fluorescent peak at the cleavage furrow is decreased.

**FIG 7.**
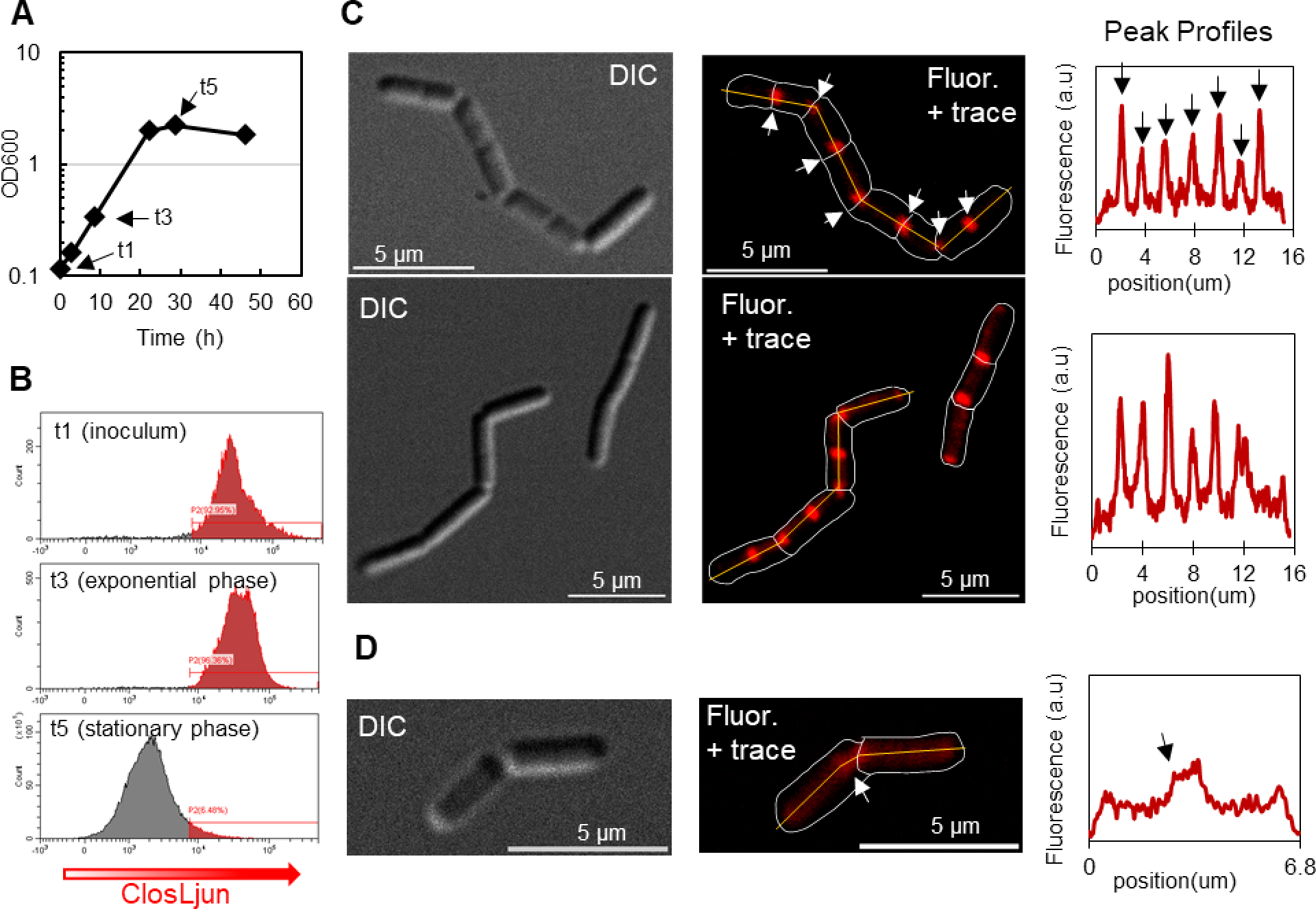
rRNA localization and ‘chaining’ in exponentially growing *C. ljungdahlii*. **(A)** The OD_600_ and **(B)** Flow Cytometric data demonstrating a steep drop off in fluorescence once the cells enter stationary phase. **(C)** DIC images of a typical *C. ljungdahlii* at early stationary phase (t3) which clearly show distinct cells bodies which have not undergone cleavage. Trace and fluorescent images show localization at the cleavage furrow and at the mid-cell. The fluorescent profile plot of the agglomerate’s major axis is indicated by the yellow line on **(C)**. Fluorescent peaks are extremely prominent during exponential phase. **(D)** Very few cells imaged microscopically from stationary phase (t5) displayed fluorescence (Fig. S12). The brightest cell from these images cell was selected and analyzed as cells in **(C)** but had decreased peak prominence compared to exponential phase.

Ribosomal compartmentalization can be seen in *C. kluyveri* though the effect is less pronounced, and the phenomenon does not appear universally even during exponential phase. Four exponentially growing cells and their fluorescent profile are shown in Fig. 8C and Fig. S13. Cell 3 has distinct fluorescent puncta at the poles, labelled α and β. The translation machinery of the other cells (cells 1, 2, and 4) is more homogeneously distributed. Unlike *C. ljungdahlii*, *C. kluyveri* (which does not sporulate either under these culture conditions) remains fluorescent into late stationary phase (Fig. 8D and Fig. S14). This observation suggests that *C. kluyveri,* with its unique substrate requirements has the ability to maintain an active translation machinery even late in stationary phase, persisting in natural milieus to scavenge its low energy-content substrates ethanol and acetate.

**FIG 8.**
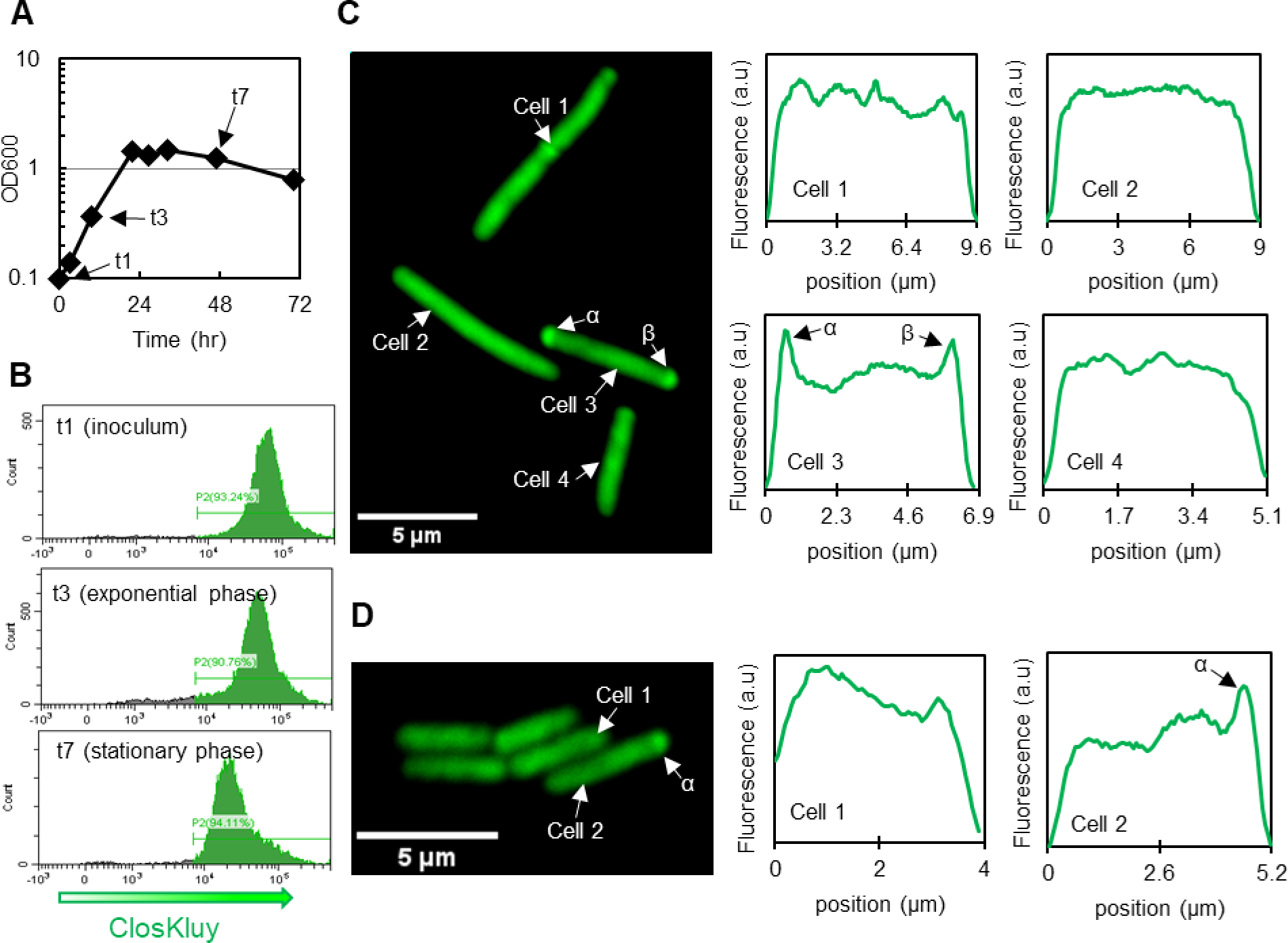
rRNA localization in C. kluyveri. **(A)** The OD_600_ and **(B)** Flow Cytometric data demonstrating a modest decrease in fluorescence once the cells leave exponential phase. **(C)** Cells exhibited ribosomal localization during exponential growth (t3) to varying extents epitomized by the 4 cells in this grouping. Cells 1, 2, and 4 show very little localization patterns, but cell 3 has distinct puncta at its poles, suggesting ribosomal localization is a transient phenomenon in *C. kluyveri* **(D)** In late stationary phase (t7), the cells remained highly fluorescent, unlike in *C. ljungdahlii,* but mostly lacked fluorescent peaks.

The surprising behavior is that of *C. acetobutylicum* (Fig. 9). Exponential phase cells (OD_600_ = ∼4) were passaged into fresh media and went quickly in exponential growth as evidenced by the straight growth line on the semi-log OD_600_ graph (Fig. 9A). As a population, the cells in the inoculum were very bright (Fig. 9B). Individually, cells exhibited a uniform distribution of translation machinery as evidenced by confocal microscopy and the accompanying fluorescence intensity plots (Fig. 9C and Fig. S15). To varying extents, weakly fluorescent puncta and ‘banding’ can be found in some cells, but these represent a minority of cells and are not representative. Two hours after passaging, the fluorescent intensity of the cells as measured by flow cytometry had decreased sharply, giving rise to a second non-fluorescent population of presumably translation-inactive cells (Fig. 9B). Full frame images in Fig. S16 and S17 show that roughly half of cells have no fluorescent signal. This was not expected as one assumes that during exponential cell growth cells are viable and actively translating proteins, though there appears to be a larger discrepancy between CFU/ml and cell counts at early exponential phase than at mid-exponential phase (33) This is a novel observation and perhaps unique among the three *Clostridium* species examined here. Among fluorescent cells, the ribosomes had localized to the poles and the mid-cell region (Fig. 9D, Fig. S16, S17). During early stationary phase, the cells became appreciably brighter with more uniformly distributed fluorescence, resembling cells in the inoculum, though the most prominent peaks were still located at the poles and some banding can be seen (Fig. 9E, Fig. S18).

**FIG 9.**
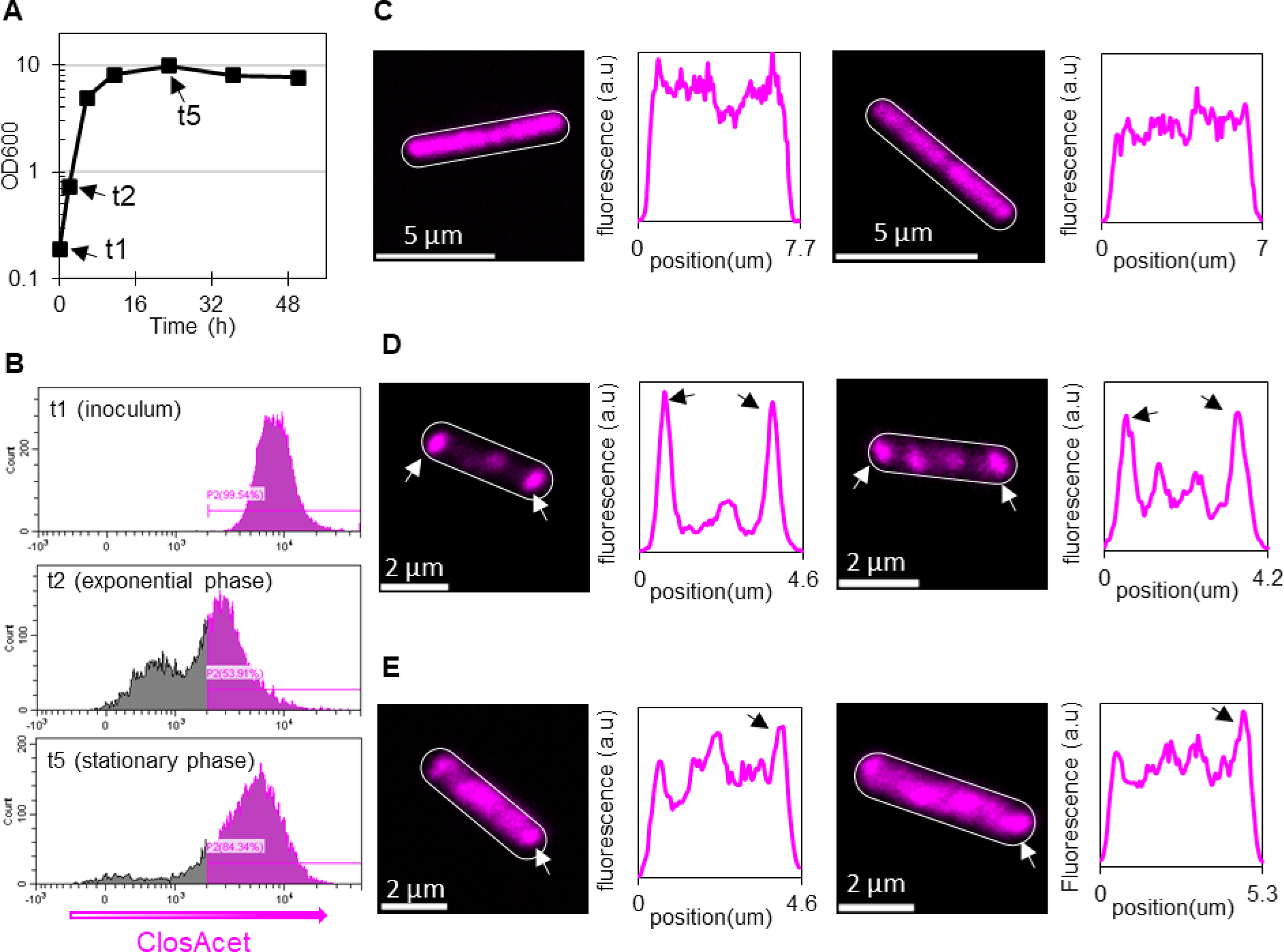
rRNA localization in *C. acetobutylicum*. An inoculum grown from a colony to an OD_600_ of ∼ 4 Our rRNA-FISH method was used to analyze the inoculum (t1), exponential (t2), and stationary phase (t5) of the culture. **(A)** The OD_600_ **(B)** Flow cytometry histograms of fluorescence in the population show a decrease in overall fluorescence during early exponential phase, but sustained fluorescence in early stationary phase. **(C)** Microscopy images and fluorescent profile plots of typical inoculum cells show no localization. **(D)** Microscopy images and fluorescent profiles plots of typical exponential phase cells indicate strong ribosomal localization. **(E)** Microscopy images and fluorescent profile plots of *C. acetobutylicum* in early stationary phase.

## DISCUSSION

Even though rRNA-FISH has been applied to prokaryotic biology for over thirty years, the development of *species-specific* rRNA-FISH probes in *clostridium spp.* is somewhat rare. We attempted to use probeBase to determine the extent to which *species-specific* rRNA-FISH probes had been designed in *clostridium spp.* (43), but the list seems incomplete since it doesn’t include some of the probes listed from other sources (12). We were able to find probes designed to target *C. leptum* (44, 45), *C. difficile* (46, 47), and *C. perfringens* (46). We could only find two examples of species-specific probes being applied to industrially interesting *Clostridia spp.* A set of probes for *C. kluyveri*, “KCLZ,” was designed to identify the species in samples from “pit mud” in the Luzhou liquor manufacturing facility (48). As previously mentioned, ClosKluy and ClosCarb had been recently designed to analyze *C. kluyveri* and *C. carboxidivorans*, respectively (18).

rRNA-FISH is suitable for rapid adoption in the field of synthetic consortia microbiology owing to its accessibility and applicability. From our experience, the PROBE_DESIGN tool in ARB is straightforward and can be used for free. The probes are easily synthesized through commercial vendors and at low-cost. Additionally, a wide array of fluorophores is available allowing greater flexibility in experimental design. In our case it was important to select probes which could be hybridized under similar conditions to allow for multiplexing, so we selected probes with similar melting temperatures as determined by freely available tools such as “OligoAnalyzer.” In addition to cytoplasmic exchange studies, rRNA-FISH has several advantages over other methods as a tool for subpopulation tracking [Fig. 4-6 and Refs. (27, 49)] as well as for identifying the fraction of actively translating (and thus actively metabolizing) cells, which, as we have shown here (Fig. 7-9) is very different for different species. Fluorescent reporter proteins could be used to identify cells but require genetic modifications, which are not available or practical in many interesting strains such as *C. kluyveri*. Dyes could be used to label the populations initially but are diluted as the cells grow. qPCR has been used to quantify sub-populations in other co-cultures (5), but provides no information about the health or expansion potential of the populations and is tedious when temporal population profiles are desirable. Modelling frameworks aiming to predict subpopulation dynamics mathematically have been proposed (50–53). So far, these approaches are only applicable in a qualitative sense, so the collection of reliable and quantitative experimental data is paramount.

We explored cytoplasmic material exchange via rRNA-FISH probes between *C. acetobutylicum*, *C. ljungdahlii* and *C. kluyveri* identifying for the first-time cellular exchange (rRNA-evidence based hybrids) between *C. ljungdahlii* and *C. kluyveri* (Fig. 4) and also strengthening the evidence for *C. acetobutylicum* - *C. kluyveri* and *C. acetobutylicum* - *C. ljungdahlii* hybrids. The rRNA hybrids that arose in the *C. kluyveri* - *C. ljungdahlii* co-cultures had a large degree of physiological heterogeneity, sometimes resembling *C. ljungdahlii*, sometimes resembling *C. kluyveri*, and sometimes resembling neither. It was essential to perform the ribosomal localization studies of each species in monoculture to better interpret the physiology of hybrid cells. Information regarding the typical, parental physiology of cells involved in cytoplasm exchange can be combined with fluorescent reporters (7, 9). The physiology of cells in Fig. 4E are characteristic of *C. ljungdahlii.* Under our experimental conditions, ‘chaining’ was only observed in *C. ljungdahlii* (Fig. 7), and that is apparently retained in some of the *C. ljungdahlii-C. kluyveri* hybrid cells (Fig. 4E). Additionally, the localization pattern of ClosKluy and ClosLjun labelled rRNA resemble *C. ljungdahlii*. Distinct puncta are associated with the nascent cleavage furrow between cells (Fig. 7C and Fig. 4E).

It is our impression that the rate at which researchers are establishing new combinatorial examples of synthetic consortia is beginning to outpace our knowledge of their fundamental biological dynamics. Moreover, if this exchange behavior, especially genetic exchange, is widespread in nature, then it would greatly affect our current understanding of evolutionary ecology in clostridial microbiomes. While this work focuses on industrially interesting *clostridia spp.*, the most conspicuous strains are pathogenic. *Clostridium perfringens* and *Clostridium septicum* are histotoxic species responsible for gas gangrene in humans and livestock (54). *C. difficile* is the leading cause of infective diarrhea in hospitals and therefor presents a tremendous burden on the healthcare systems worldwide (55). Additionally, botulism and tetanus are caused by clostridial infections. Understanding interspecies relationships among these species would provide far-reaching insights into agriculture and human health, as well as industrial fermentation (56).

## MATERIALS AND METHODS

### Probe Design

All probes used in this study can be found in Table S1. The ClosAcet and ClosLjun probe sets were designed and named similarly to Schneider et al., who demonstrated an analogous system on *C. carboxidivorans* and *C. kluyveri* (*18*). Briefly, the PROBE_DESIGN tool in ARB (Version 7.0) was used with the SILVA LSU Ref NR 99 (version 138.1) library to generate potential probe sequences (https://www.arb-silva.de/). ClosAcet was designed to target all entries within the *C. acetobutylicum* ATCC 824, DSM 1731, EA2018 subtree. Designing a probe which labelled only *C. ljungdahlii* (and not the closely related *C. autoethanogenum*) was deemed impossible, since the only suitable probe coincidentally had specificity for *C. acetobutylicum*. Therefor ClosLjun was designed with specificity to *C. autoethanogenum* DSM 10061 and *C. ljungdahlii* DSM 13528. Probe selections were made based on the ARB’s estimated selectivity of the probe (i.e. ‘eqaul’). We intended to label multiple species simultaneously, so it was essential that the ideal hybridization stringency (determined by formamide concentration in the hybridization buffer) be similar between all probe species. We hypothesized that probes with similar melting temperatures would share optimal hybridization conditions. ARB provides a basic melting temp based on probe length and GC content, but nearest-neighbor (NN) methods are far more accurate and parameters specifically for RNA/DNA duplex formation are readily available (57). We used the “OligoAnalyzer” tool from Integrated DNA Technologies (IDT) to select probes with melting temperatures between 68-72 °C presuming a 1 μM probe concentration and 900 mM Na+ concentration. (https://www.idtdna.com/calc/analyzer). Cy3 was chosen as the fluorescent tag for ClosAcet and Cy5.5 was chosen as the fluorescent tag for ClosLjun. These fluorescent molecules are known to be orthogonal green fluorophores (such as Alexa Fluor 488; used with ClosKluy and the OregonGreen HaloTag Ligand) on the fluorescent channels used for flow cytometry and microscopy. Orthogonality was checked with the fluorescent spectrum tool from AAT BioQuest. (https://www.aatbio.com/fluorescence-excitation-emission-spectrum-graph-viewer). All probes were synthesized by IDT.

### Microorganisms and growth media

Monocultures of *C. acetobutylicum* (ATCC824), *C. ljungdahlii* (ATCC 55383), and *C. kluyveri* (ATCC 8527) were grown in Turbo Clostridial Growth Medium (TCGM) supplemented with species-specific carbon sources and nutrients as previously described (5, 6). The TCGM base stock contains the following amounts of each component per liter: yeast extract, 5 g; asparagine, 2 g; 50x mineral stock solution, 20 mL; 100x trace elements solution, 10 mL; phosphate buffer solution, 10 mL; 100x Wolfe’s vitamins, 10 mL; 500x PABA, 2 mL. The 50x mineral stock solution contains the following amount of each component per liter: NaCl, 50.5 g; MgSO_4_, 17.4 g; (NH_4_)_2_SO_4_, 100 g; sodium acetate, 123 g. The 100x trace elements solution contains the following amount of each component per liter: Nitrilotriacetic acid, 1.5 g; MnSO_4_*7H_2_O, 1.5 g; FeSO_4_*7H_2_O, 1.1 g; CoCl_2_*6H_2_O, .1 g; CaCl_2_, 2.1 g; ZnSO_4_*7H_2_O, .18 g; CuSO_4_*5H_2_O, 10mg; KAl(SO_4_)_2_*12H_2_O, 20 mg; H_3_BO_3_, 10 mg; Na_2_MoO_4_*2H_2_O, 10 mg; NiCl_2_*6H_2_O, 30 mg; Na_2_WO_4_*2H_2_O, 20 mg; Na_2_SeO_3_*5H_2_O, 300 μg. 100x Wolfe’s vitamins contains the following amounts per liter: pyridoxine-HCl, 10 mg; thiamine-HCl, 5 mg; riboflavin, 5 mg; Calcium pantothenate, 5 mg; thioctic acid, 5 mg; para-aminobenzoic acid, 5 mg; nicotinic acid, 5 mg; Vitamin B_12_, 100 μg; d-biotin, 2 mg; folic acid, 2 mg. The phosphate buffer contains the following amounts per liter: KH_2_PO_4_, 100 g; K_2_HPO_4_, 125 g. The phosphate buffer is adjusted to a pH of 6.9 with NaOH. The 500x PABA solution contains 2 g/L para-aminobenzoic acid. The 100x phosphate buffer, 100x Wolfe’s vitamins, and 500x PABA are filter sterilized. Yeast extract, asparagine, 100x trace elements and 50x mineral solution are dissolved in 800 mL RO-H_2_O and autoclaved at 121°C for 30 min. After cooling, 100x phosphate buffer solution, 100x Wolfe’s vitamins and 500x PABA are added. For cultures containing *C. acetobutylicum*, TCGM was supplemented with 160 mL/L of a filter-sterilized 500 g/L glucose solution and 10 mL/L of a filter sterilized 500 g/L fructose solution (80/5 TCGM). For cultures containing *C. ljungdahlii*, TCGM was supplemented with10 mL/L of filter sterilized 500g/L fructose (0/5 TCGM). For cultures containing *C. kluyveri*, TCGM was supplemented with 20 mL/L ethanol, 40 mL/L of 200 g/L sodium acetate solution, 25 mL/L of a 100 g/L sodium bicarbonate, 10 mL/L of a 15 g/L L-Cysteine-HCl solution, and an additional 10 mL/L of the 100x phosphate buffer (TCGM-Ckl). The balance was sterile RO-water. All media were left to deoxygenate in an anaerobic chamber (Forma Anaerobic System; Thermo Fisher Scientific) for at least 2 days. For growth of solid media, 2xYTG plates were used which contain the following per liter: tryptone, 16 g; yeast extract, 10 g; NaCl, 4 g; glucose, 5g; agar, 15g. After combining all the ingredients, the media is adjusted to a pH of 5.8 with 5M HCl before autoclaving. The media is cooled to 65°C before addition of antibiotics and poured into plates immediately afterwards.

### Growth of monocultures and pre-cultures

Exponentially growing cultures of *C. acetobutylicum, C. ljungdahlii*, and *C. kluyveri* were diluted 9:1 in pure DMSO and stored at −80°C (New Brunswick Scientific, Edison, NJ, USA). For *C. acetobutylicum,* 150 μL of frozen stocks was streaked on 2xYTG plates anaerobically and allowed to incubate at 37°C for more than 4 days to allow for sporulation. After this, plates were stored aerobically at 4°C for up to a year to select against vegetative cells which would be killed in exposure to oxygen. To begin *C. acetobutylicum* liquid cultures, 10 mL of 80/5 TCGM is inoculated with a single colony and heat shocked for 10-20 min. at 80°C to initiate spore germination. Growth was typically observed after 14-20 hours. To prevent acid death, the culture’s pH is adjusted to 5.2 with NaOH after 22-28 hours, unless passaged beforehand. To begin the *C. ljungdahlii* liquid cultures, 1 mL frozen stocks of *C. ljungdahlii* were sowed anaerobically into 9 mL of 0/5 TCGM and passaged as needed after 16 hours once the cell reached log-phase. For *C. kluyveri*, 1 mL of cells were sowed into 19 mL of TCGM-Ckl. The plasmid of *C. acetobutylicum*-p100ptaHALO and *C. ljungdahlii*-p100ptaHALO was maintained by addition of 100 μg erythromycin (Em) per mL of culture media from a 1000x concentrated stock. When growing *C. acetobutylicum* from colonies, Em was added after heat shock.

### C. kluyveri and C. ljungdahlii co-culture preparation

A special culture medium was prepared for the *C. kluyveri* and *C. ljungdahlii* co-culture by omitting acetate entirely from TCGM-Ckl and adding 4 mL/L of the 500 g/L fructose solution (TCGM-Ckl/Clj). 20 mL precultures of *C. ljungdahlii* and *C. kluyveri* were grown to exponential phase (OD_600_ of .4 to .6) in 0/5 TCGM and TCGM-Ckl, respectively. To initiate the co-culture, 10 mL of each preculture was separately washed once in TCGM-Ckl/Clj and resuspended into 1 mL. The two 1 mL cell concentrations are then added to 13 mL of fresh TCGM-Ckl/Clj in a sealed Balch tube (18mm x 150mm, ChemGlass), resulting in a 15 mL co-culture with an R value of 1. The headspace of the Balch tube was filled to between 35 and 45 psi of an 80/20 blend of H_2_ and CO_2_ gas. After each sampling, the head space was repressurized.

### C. acetobutylicum and C. ljungdahlii Co-culture preparation

A 90 mL culture of *C. ljungdahlii*-p100ptaHalo was grown to exponential phase (OD_600_ of .4 to .6) and a 30 mL culture of *C. acetobutylicum* was grown to early stationary phase (OD_600_ of 6 to 8). The full 90 mL *C. ljungdahlii*-p100ptaHalo culture was washed three times to remove residual erythromycin and concentrated to 1 mL. 5 mL of *C. acetobutylicum* was washed and resuspended in 29 mL of 80/5 TCGM. The high-density *C. ljungdahlii*-p100ptaHalo inoculum was added to the *C. acetobutylicum* to start the co-culture (5).

### C. acetobutylicum, C. ljungdahlii, C. kluyveri triple co-culture preparation

To prepare the triple co-culture medium, TCGM-Ckl was supplemented with 3.5 g/L glucose and 3 g/L fructose. *C. ljungdahlii* and *C. kluyveri* were grown separately to early exponential phase (OD_600_ of .2 to .4) in 40 mL cultures, and *C. acetobutylicum* was grown to early stationary phase (OD_600_ of 6 to 8) in a 10 mL culture. A volume of cells corresponding to 4 OD_eq_ (where OD_eq_ is the product of the volume (mL) and the OD_600_) of each species is spun down and washed in the triple culture media twice and resuspended in 3.33 mL. Then the samples are combined to form a 10 mL triple culture.

#### *In-solution* RNA-FISH protocol

Culture samples were washed twice in filter-sterilized ice-cold 1xPBS (Gibco, pH 7.4) via centrifugation (5-10 min, 3220 rcf, 4°C, Centrifuge 5810 R; Eppendorf). Then the pellet was resuspended thoroughly in ice-cold 1xPBS and fixed by 1:1 dilution in ice-cold absolute ethanol. All samples are stored at −20°C for up to a month.

For each *in-solution* RNA-FISH sample, 0.15 OD_eq_ of cells were pelleted via centrifugation 10 min., 10000 rcf, 4°C, Z216 MK; Hermle) in a 1.6 mL microcentrifuge tube. The OD_eq_ is calculated by the following equation:

OD_eq_ = OD_600_*Volume (mL)

After carefully removing the supernatant, the pellet was dried for 5 to 15 minutes at 46°C (Isotemp Hybridization Incubator; Fisher Scientific) to remove residual ethanol. The pellet was resuspended in 75 μL of hybridization buffer containing 0.9 M NaCl, 0.02 M Tris-HCl (pH 7.0), 20% (v/v) formamide, 0.01% sodium dodecyl sulfate, and 1 μM probe(s) in aqueous solution and incubated at 46°C for 5 hours, unless otherwise noted. During hybridization 50 mL of wash buffer was prepared containing 0.215 M NaCl, 5 μM EDTA, 0.02 M Tris-HCl (pH 7.0), and 0.01% sodium dodecyl sulfate and pre-warmed to 48°C. Immediately following hybridization, the cells are pelleted via centrifugation (8-10 min., 10,000 rcf, Centrifuge 5418; Eppendorf), and the supernatant is discarded into as formamide waste stream. The cells are washed twice via incubation in 500 μL of washing buffer for 20 min at 48°C. Finally, the cells are washed once and resuspended in 1 mL of ice-cold 1xPBS. For the composition of hybridization and washing buffers used throughout the manuscript, refer to Table S2.

### Deep Red Labelling

CellTracker Deep Red was applied as previously described (9). Briefly, 15μg of CellTracker Deep Red was dissolved in 20μL of dimethyl sulfoxide (DMSO) to generate an 1000x (1 mM) labelling stock. The 1000x stock was added directly to the culture media and incubated at 37°C for 40 min. Excess dye was removed by washing the cells three times in medium via centrifugation and resuspension.

### HaloTag labelling

The HaloTag protein was labelled as previously described (31). Prior to fixing, 0.5 μL aliquots of concentrated ligand were added to 500 μL aliquots of cells and incubated at 37°C for 15-60 minutes anaerobically. The final ligand concentration for Janelia 549 and 646 was 200 nM. The final ligand concentration for OregonGreen was 1 μM. After labelling, the cells were washed twice in warm 1xPBS. Cells labelled with OregonGreen were further resuspended in fresh, warm culture media and incubated for 30 min. at 37°C to wash out any residual ligand. After one final wash in 1xPBS, the cells were ready for flow cytometry, microscopy, or pre-hybridization fixing.

### Flow cytometry

A flow cytometer (CytoFLEX S, Beckman Coulter Life Sciences, Indianapolis, IN, USA) was used to collect fluorescence and light scattering data from individual cells. CytExpert (Beckman Coulter, Version 2.4.0.28) was used as the acquisition and data processing software. Cells were diluted to approximately 10^5^ cells per μL, (c.a. OD_600_ of .1) in 1xPBS on ice. A 488 nm laser was used to collect forward angle light scatter (FSC) and right-angle side scatter (SSC) for the sample. Only events surpassing a pulse height of 900 on the SSC channel were counted. The sample feed rate was adjusted between 10 and 60 μL/min to reach a sampling rate of about 1,000 events/sec. In all samples, the abort rate was kept below 7% by adjusting sampling rate. For each sample, 5 μL was interrogated. The fluorescent signal of ClosKluy and OregonGreen was excited by a blue laser and filtered through a 525/40 band pass filter (BPF) before acquisition. The fluorescent signal of ClosAcet and Jenalia 549 was excited by yellow laser and filtered through a 585/42 BPF before acquisition. The fluorescent signal of ClosLjun, Jenalia 646, and Deep Red was excited by a red laser and filtered through a 712/25 BPF before acquisition. The gain of the avalanche photodiodes was adjusted to 125, 1000, and 850 to acquire ClosKluy, ClosAcet, and ClosLjun, respectively. The standard gain following QC was used for the HaloTag ligands and Deep Red. For each channel, the fluorescent intensity is given by the area under the curve of emission intensity, rather than height. For each sample, the median of the population was used to determine central tendency.

### Microscopy and Image analysis

Following *in-solution* rRNA-FISH, between 5 and 50 μL of cell sample was dropped onto either poly-l-lysine coated chambered slides (Nunc Lab-Tek 8-Well chambered slides, Thermo scientific) or 10-well diagnostic slides (MP Biosciences, Irvine, CA, USA) and dried completely at 46°C to ensure adhesion. Cells in the chambered slides were submerged in 200 μL of cold 1xPBS prior to imaging. Diagnostic slides were rinsed in dunk tanks of cold RO-H_2_O, then dried quickly under compressed air to remove residual water before being interred in PVA with DABCO (Sigma Aldrich, Darmstadt, Germany). All microscopy was performed on an ANDOR Dragonfly spinning disk confocal microscope under a 63x or 100x oil objective. The fluorescent signal of ClosKluy and OregonGreen was excited by a blue laser (λ_ex_ = 488nm) and filtered through an 521/38 band pass filter (BPF) before acquisition. The fluorescent signal of ClosAcet and Jenalia 549 was excited by yellow laser (λ_ex_ = 561 nm) and filtered through an 594/43 BPF before acquisition. The fluorescent signal of ClosLjun, Jenalia 646, and Deep Red was excited by a red laser (λ_ex_ = 638 nm) and filtered through an 685/47 BPF before acquisition. Image J (https://imagej.net/ij/) was used to process microscopy images. The brightness and contrast setting were adjusted for the control images (Fig. S1, S2, and S3) to maximize signal representation. In other words, the white point was minimized to capture any signal over background. For all other images, only the white point of the image was increased to minimize oversaturation in images of very bright samples. The black point and microscope power settings were never changed.

## SUPPLEMENTAL MATERIAL

Supplemental material is available online only.

**Table S1** DOCX File

**Table S2** DOCX File

**Fig. S1** DOCX File

**Fig. S2** DOCX File

**Fig. S3** DOCX File

**Fig. S4** DOCX File

**Fig. S5** DOCX File

**Fig. S6** DOCX File

**Fig. S7** DOCX File

**Fig. S8** DOCX File

**Fig. S9** DOCX File

**Fig. S10** DOCX File

**Fig. S11** DOCX File

**Fig. S12** DOCX File

**Fig. S13** DOCX File

**Fig. S14** DOCX File

**Fig. S15** DOCX File

**Fig. S16** DOCX File

**Fig. S17** DOCX File

**Fig. S18** DOCX File

## Acknowledgements

Supported by an ARPA-E project under contract AR0001505. J.D.H. was supported in part by a U.S. Department of Education GAANN Fellowship under grant P200A210065. Microscopy equipment was acquired with a shared instrumentation grant (grant S10 OD016361), and access was supported by the NIH-NIGMS (grant P20 GM103446), by the NSF (grant IIA-1301765) and by the State of Delaware.

E.T.P. conceived the overall project. J.D.H. conceived the approach, worked with E.T.P for the experimental design, and carried out all experiments and data analysis. J.D.H. and E.T.P. interpreted the data and wrote the manuscript.

## AUTHOR CONTRIBUTIONS

John D. Hill, Conceptualization, Investigation, Methodology, Formal analysis, Validation, Visualization, Resources, Data curation, Writing – original draft, review and editing | Eleftherios Terry Papoutsakis, Conceptualization, Data curation, Formal analysis, Funding acquisition, Investigation, Methodology, Project administration, Resources, Supervision, Writing – review and editing.

## Supplementary tables

**Table S1.**
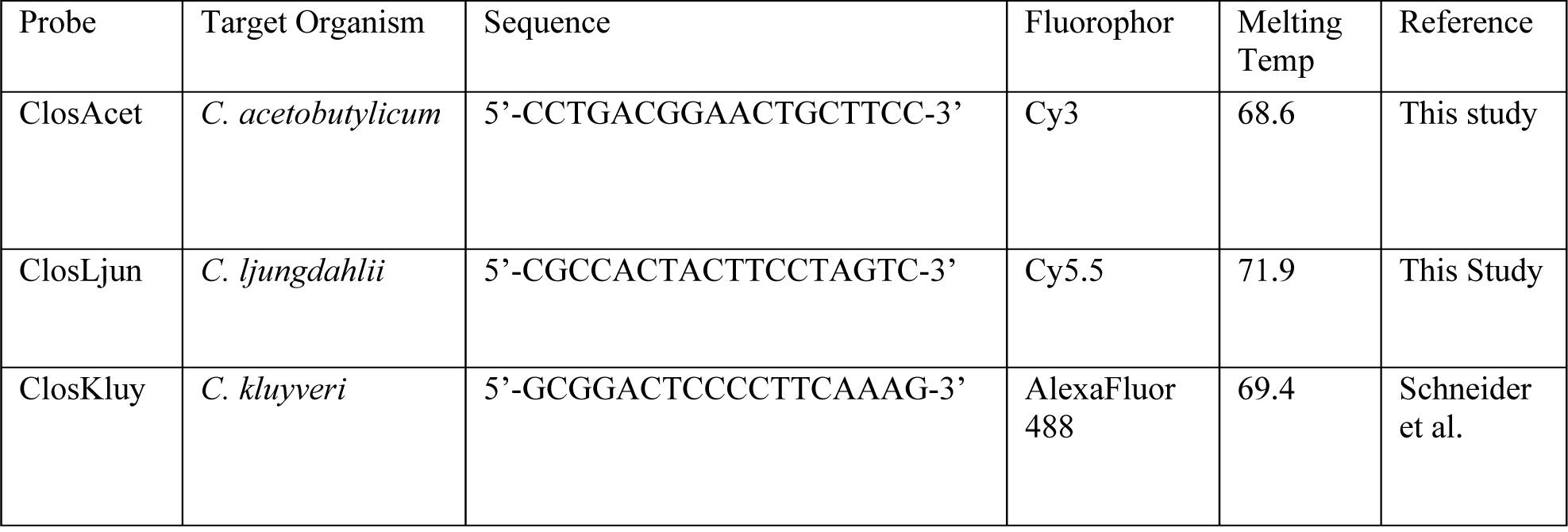
Probes used in this study.

**Table S2.** NaCl and EDTA concentration in washing buffer.

| Formamide % in Hyb. Buff. | NaCl conc. | EDTA conc. |
| --- | --- | --- |
| 10 | 0.45 M | omitted |
| 20 | 0.215 M | 5 μM |
| 30 | 0.102 M | 5 μM |
| 40 | 0.046 M | 5 μM |
| 50 | 0.018 M | 5 μM |

## Supplementary material

**Fig S1.**
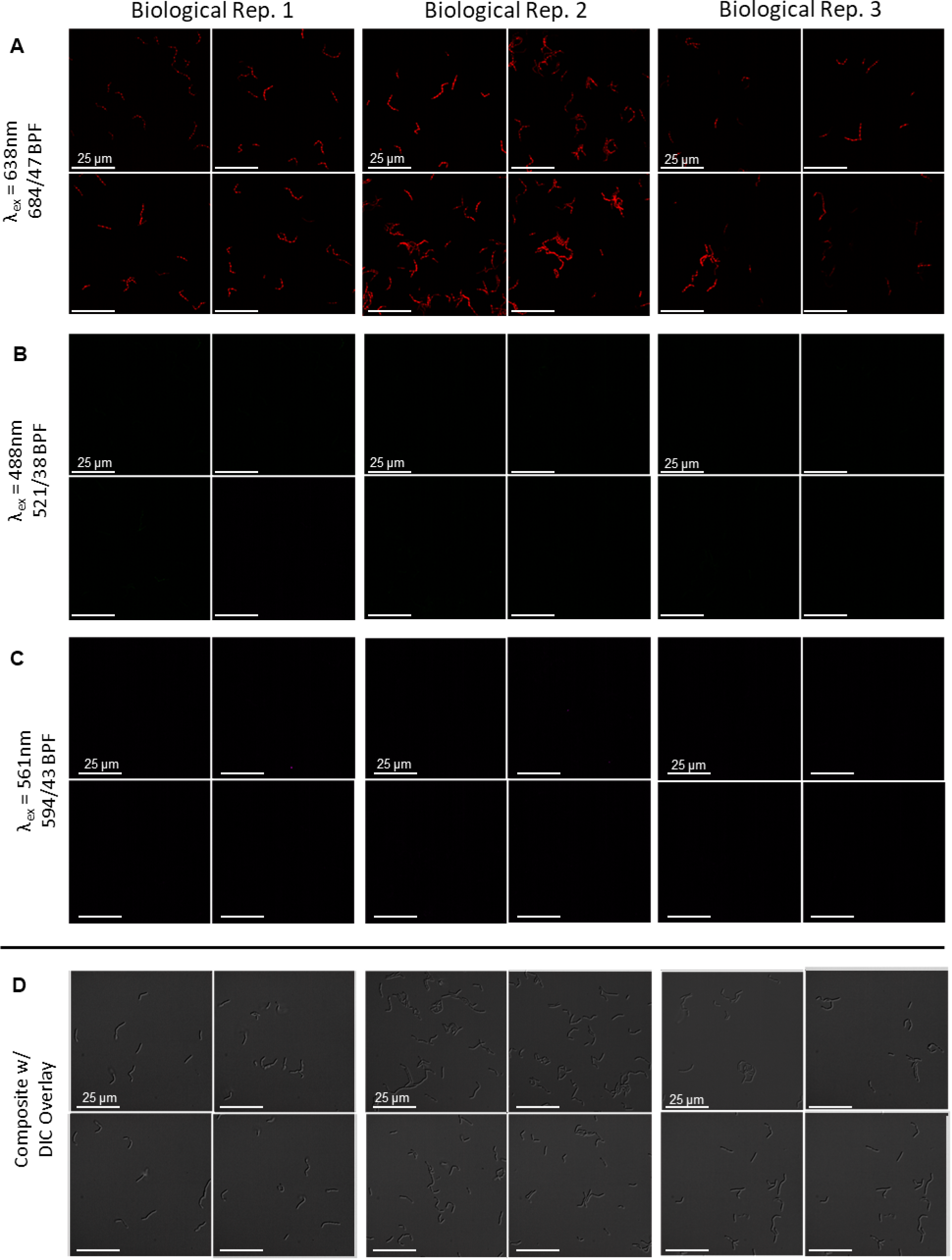
Fluorescent microscopy images of *C. ljungdahlii* from FIG 1.E. **(A-C)** 4 random fields from each of 3 labelled (incubated with all three sets of probes simultaneously) biological replicates were imaged via the far-red channel (λ_ex_=638 nm, BPF = 684/47, specific to ClosLjun) **(A),** the green channel (λ_ex_=488 nm, BPF = 521/38, specific to ClosKluy) **(B),** and the yellow channel (pseudo-colored magenta) (λ_ex_=561 nm, BPF = 594/43, specific to ClosAcet**) (C).** Only red fluorescence, coming from ClosLjun, could be detected. (**D)** Composite images (all three channels stacked) with DIC overlay reveal the absence of autofluorescence in the unlabeled cells.

**Fig S2.**
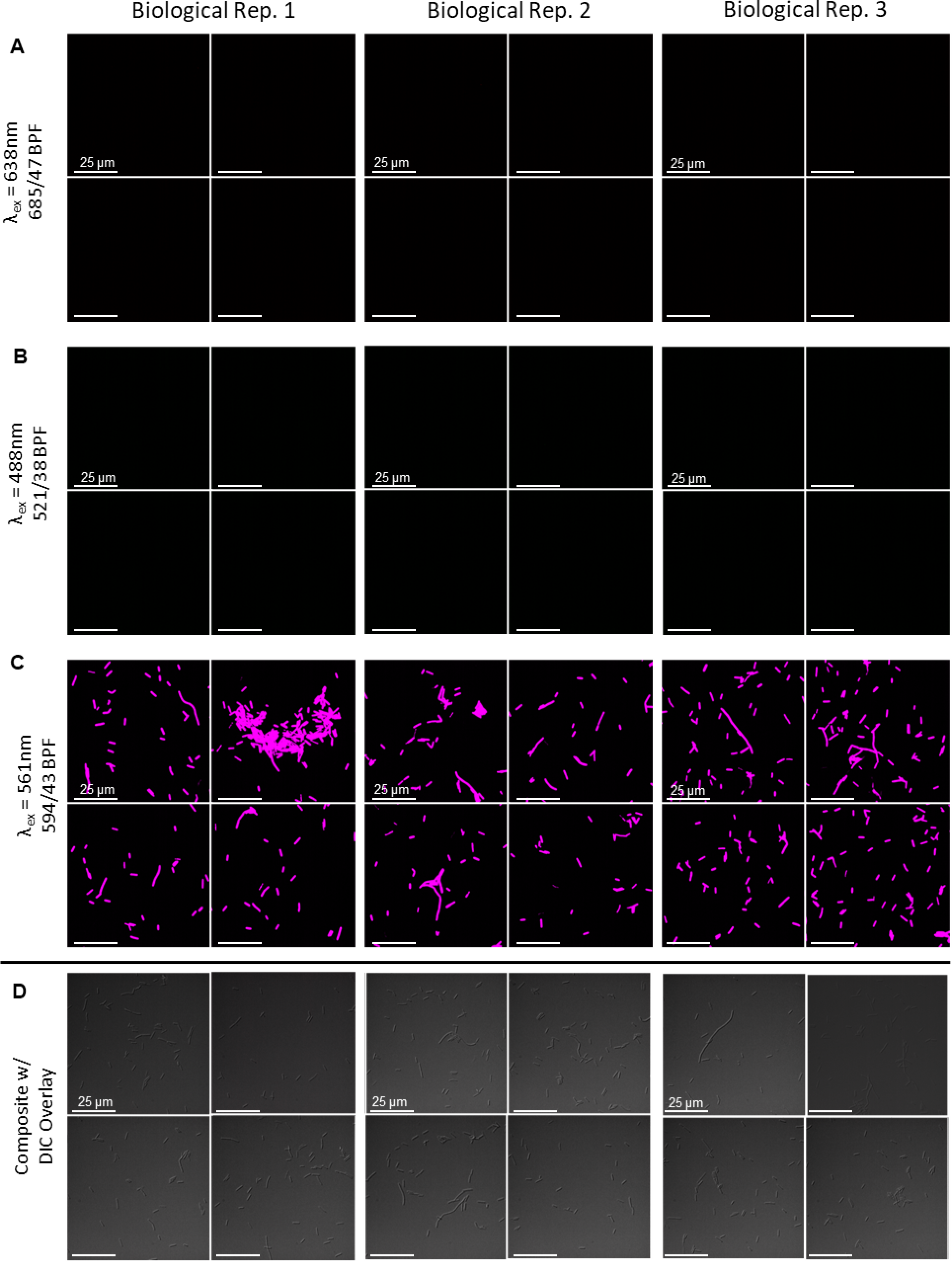
Fluorescent microscopy images of *C. acetobutylicum* from FIG 1.D. **(A-C)** 4 random fields from each of 3 labelled (incubated with all three sets of probes simultaneously) biological replicates were imaged via the far-red channel (λ_ex_=638 nm, BPF = 684/47, specific to ClosLjun) **(A),** the green channel (λ_ex_=488 nm, BPF = 521/38, specific to ClosKluy) **(B),** and the yellow channel (pseudo-colored magenta) (λ_ex_=561 nm, BPF = 594/43, specific to ClosAcet) **(C).** Only magenta fluorescence, coming from ClosAcet, could be detected. **(D)** Composite images (all three channels stacked) with DIC overlay reveal the absence of autofluorescence in the unlabeled cells.

**Fig S3.**
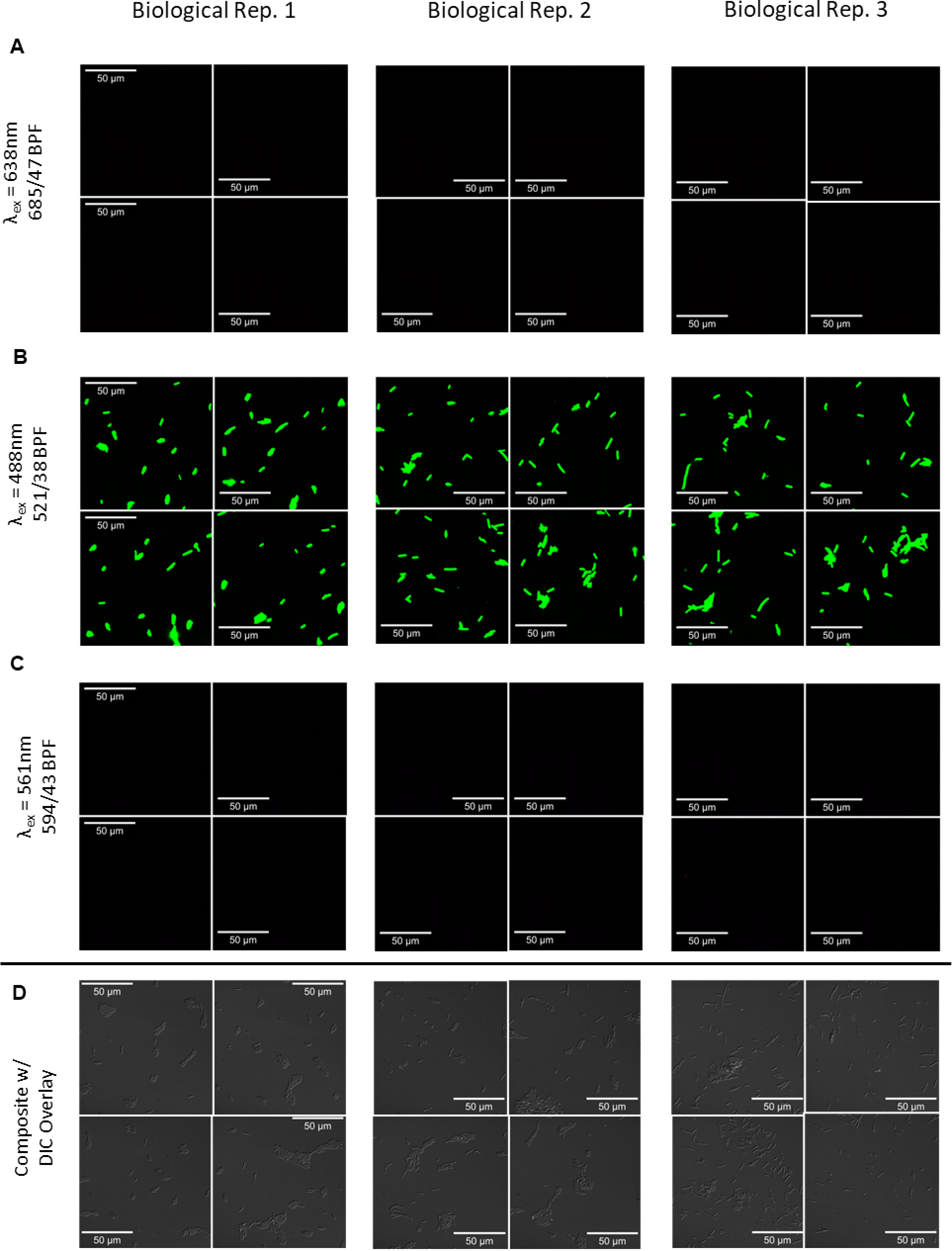
Fluorescent microscopy images of *C. kluyveri* from FIG 1.F. **(A-C)** 4 random fields from each of 3 labelled (incubated with all three sets of probes simultaneously) biological replicates were imaged via the far-red channel (λ_ex_=638 nm, BPF = 684/47, specific to ClosLjun) **(A),** the green channel (λ_ex_=488 nm, BPF = 521/38, specific to ClosKluy) (B), and the yellow channel (pseudo-colored magenta) (λ_ex_=561 nm, BPF = 594/43, specific to ClosAcet) **(C).** Only green fluorescence, coming from ClosKluy, could be detected. **(D)** Composite images (all three channels stacked) with DIC overlay reveal the absence of autofluorescence in the unlabeled cells.

**FIG S4.**
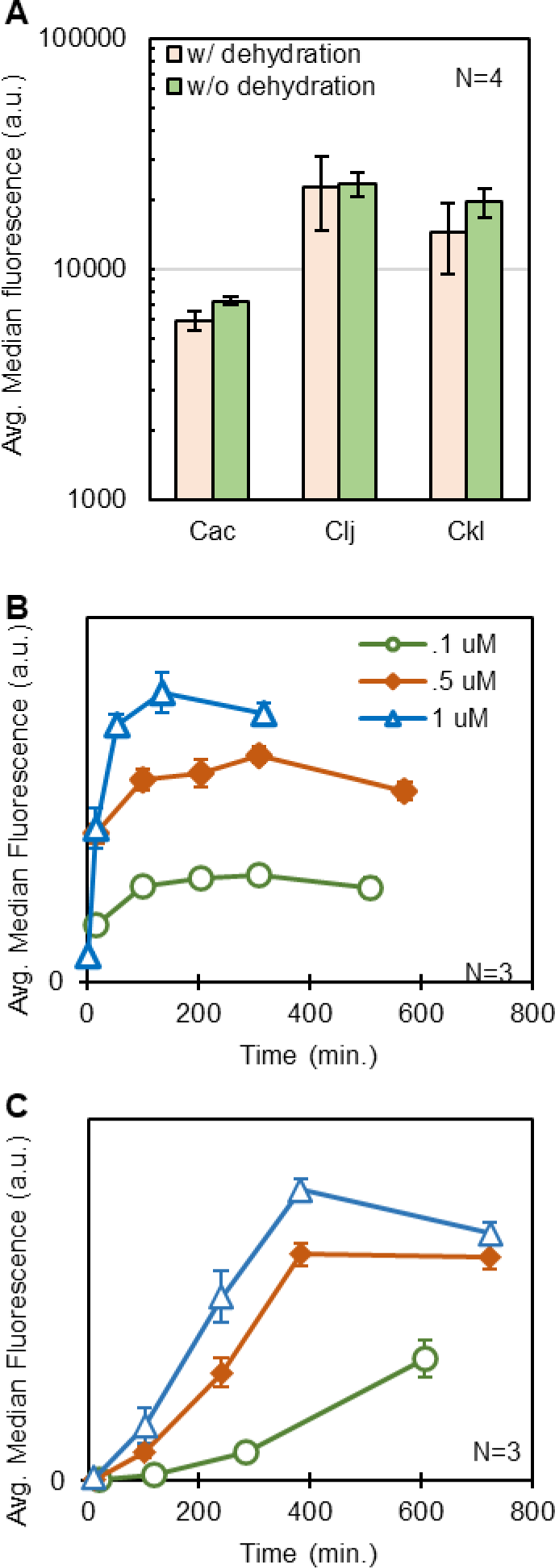
*In-solution* rRNA-FISH method optimization. **(A)** The average median fluorescence of 4 biological replicate populations of *C. acetobutylicum* (Cac), *C. ljungdahlii* (Clj), and *C. kluyveri* (Ckl) were measured with or without an additional prehybridization dehydration step. No statistically significant change was observed. Error bars represent a single standard deviation above and below the mean in all figures. **(B and C)** The probe concentration and hybridization duration were optimized. In **(B)**, 3 biological replicates of exponentially growing *C. acetobutylicum* were fixed and incubated with three different concentrations of ClosAcet. The average median fluorescence of the three replicates (in arbitrary units, a.u., linear y axis, indexed at 0) was tracked over 10 hours. **(C)** Demonstrates the analogous experiment performed in *C. ljungdahlii* with ClosLjun.

**Fig S5.**
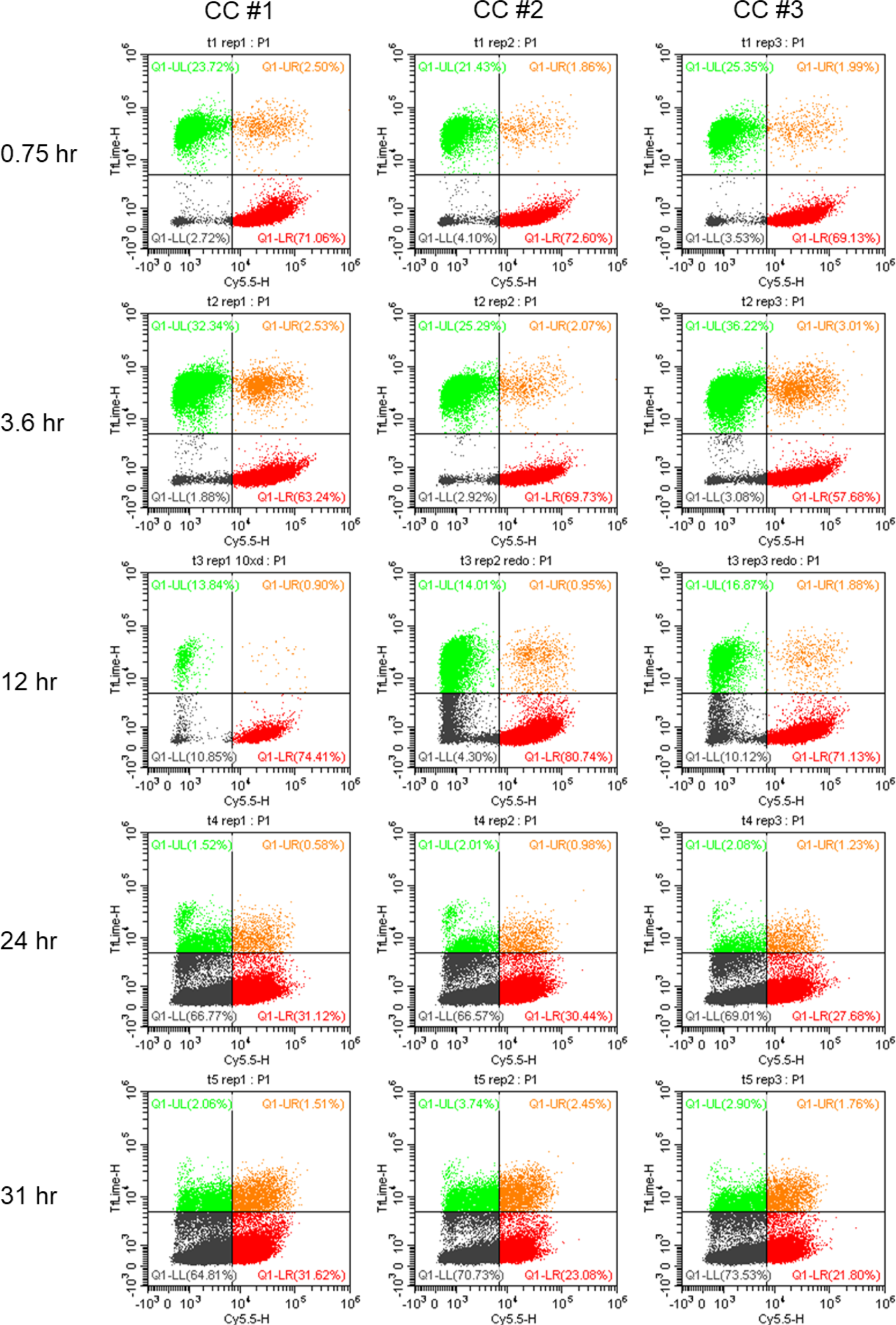
Flow cytometry dot-plots used to determine subpopulations and hybrid cell frequency in the co-culture between *C. kluyveri* and *C. ljungdahlii*.

**Fig S6.**
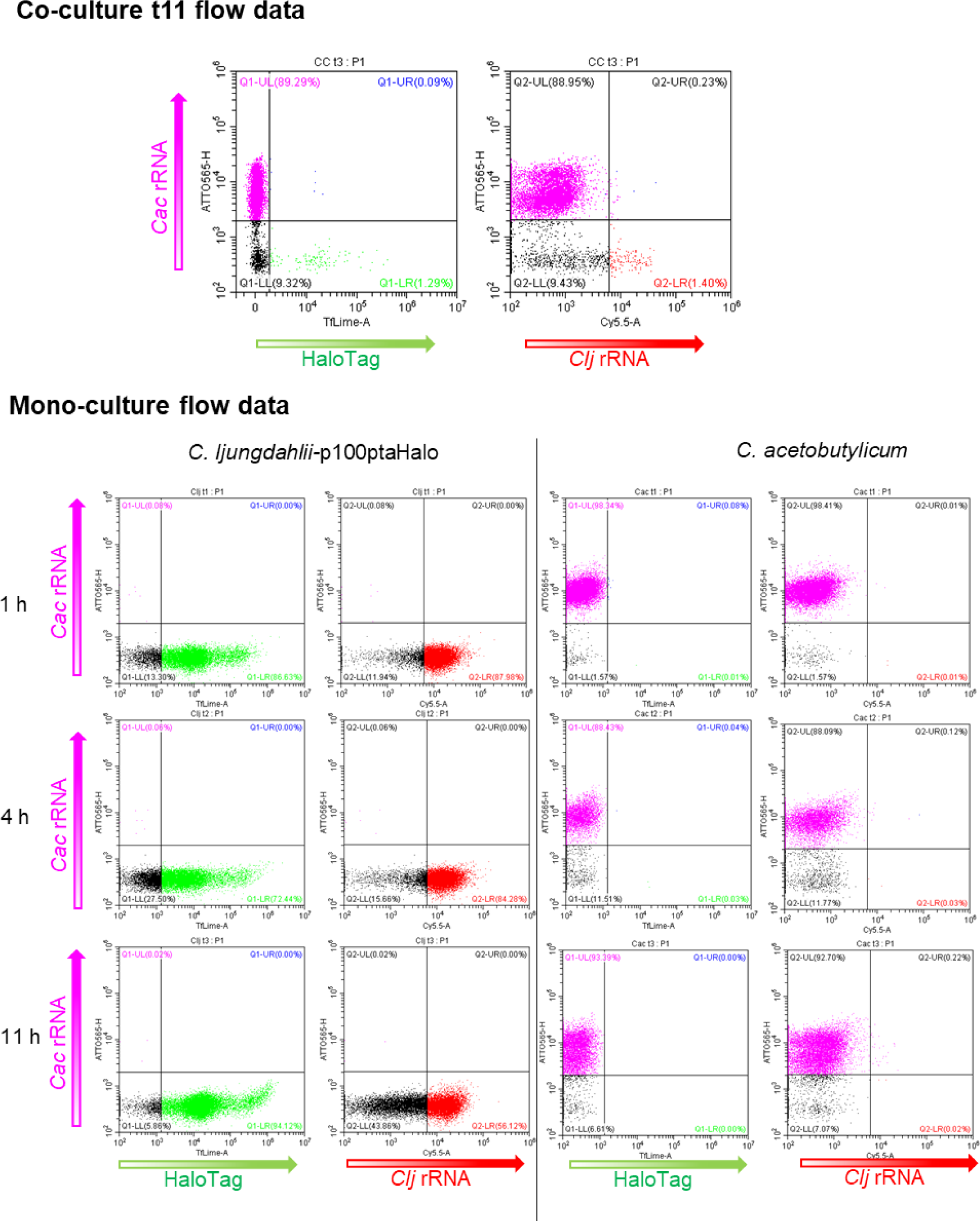
The 11-hour time point flow cytometry dot-plots and the pure-culture controls for the co-culture in Fig. 5. Double positive cells are essentially non-existent for either species across all timepoints.

**Fig S7.**
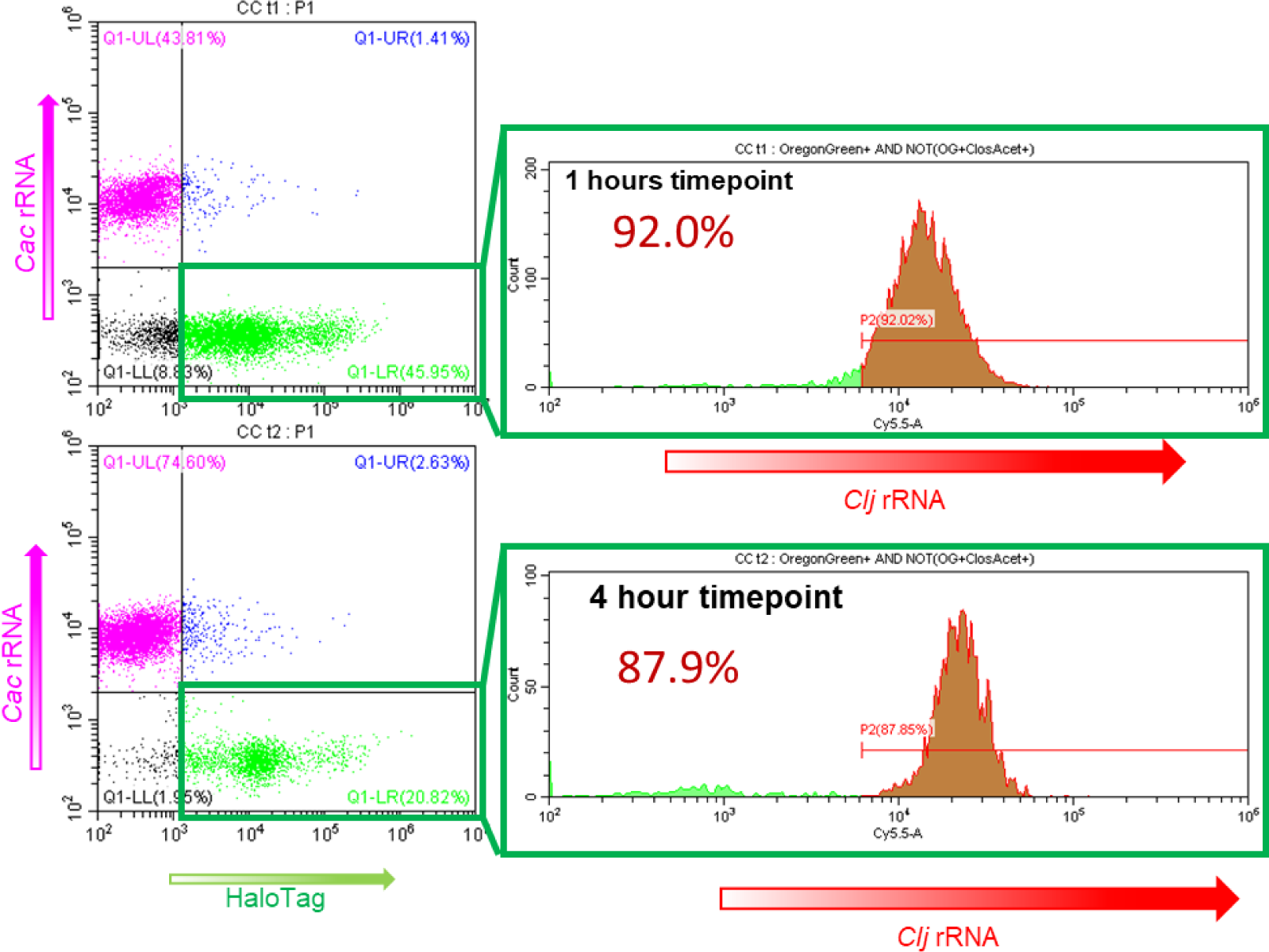
Prevalence of red fluorescent cells (i.e. high fluorescence on the ClosLjun channel) among green-fluorescent cells (excluding the hybrid subpopulation). This demonstrates that nearly all green labelled cells are double positive for the ClosLjun probe.

**Fig S8.**
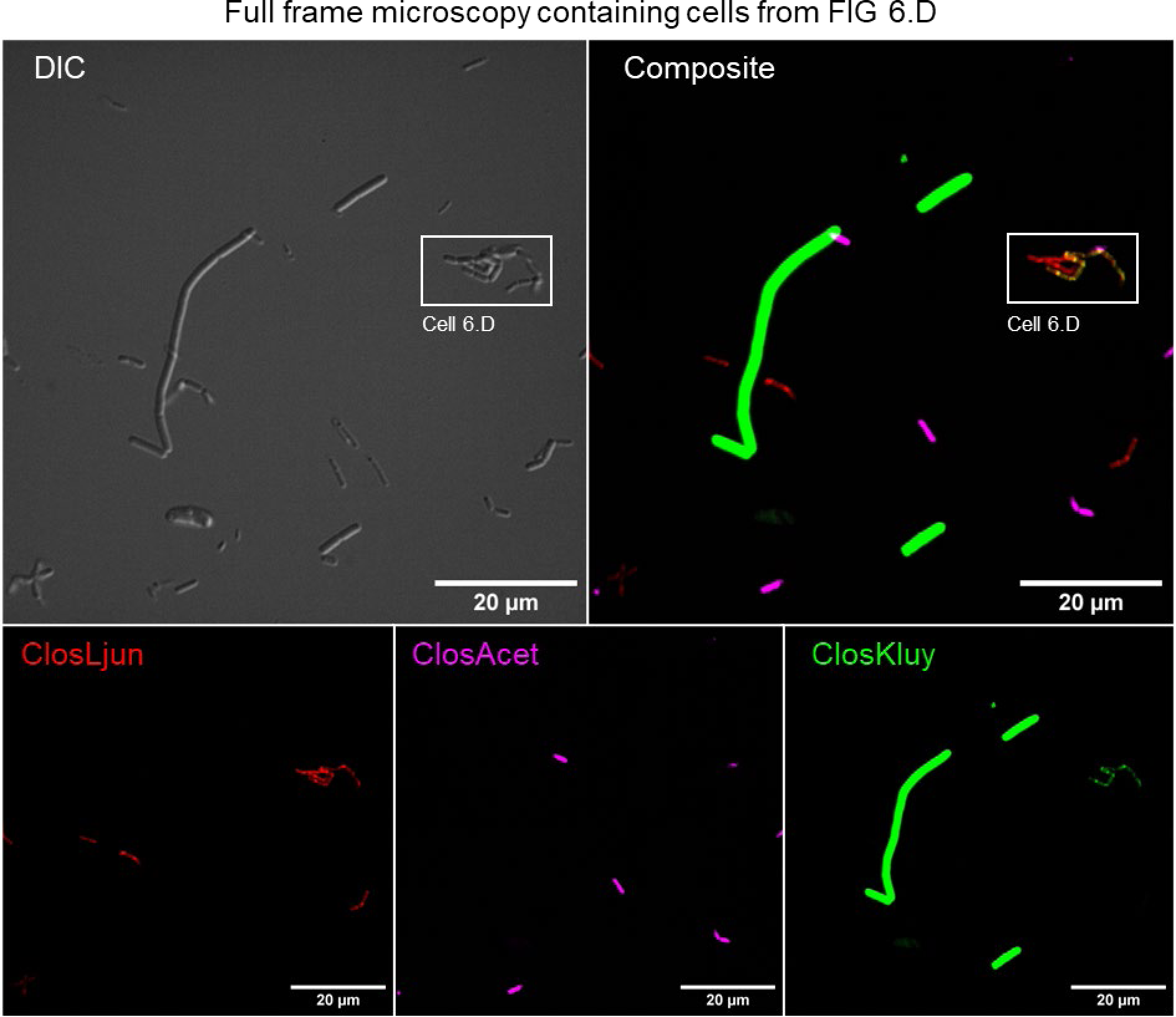
Full Frame images of the cells depicted in Fig. 6D. In the Top row, the DIC and the composite fluorescent image are presented, with the individual fluorescent channels comprising the bottom row.

**Fig S9.**
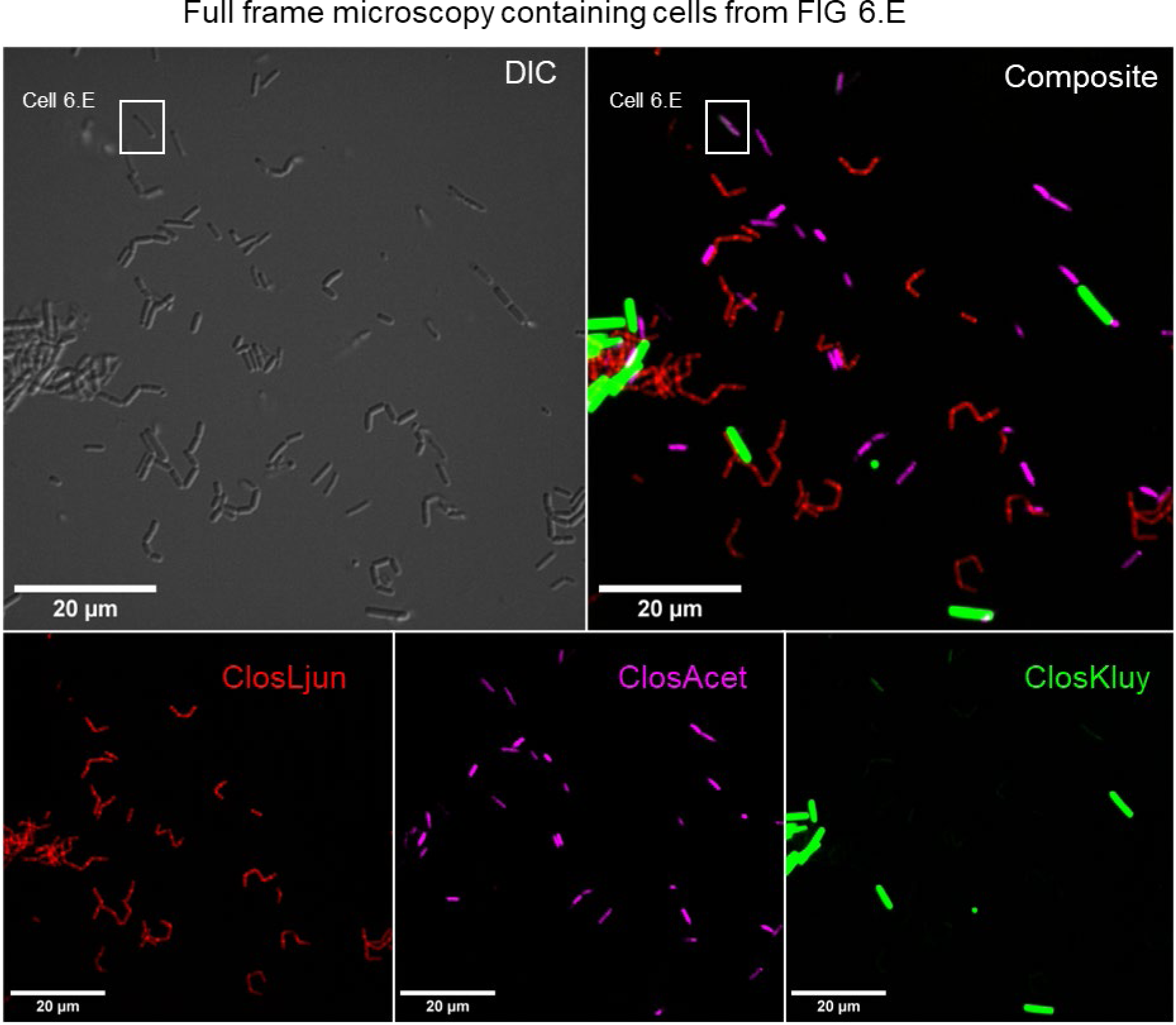
Full Frame images of the cells depicted in Fig. 6E. In the Top row, the DIC and the composite fluorescent image are presented, with the individual fluorescent channels comprising the bottom row.

**Fig S10.**
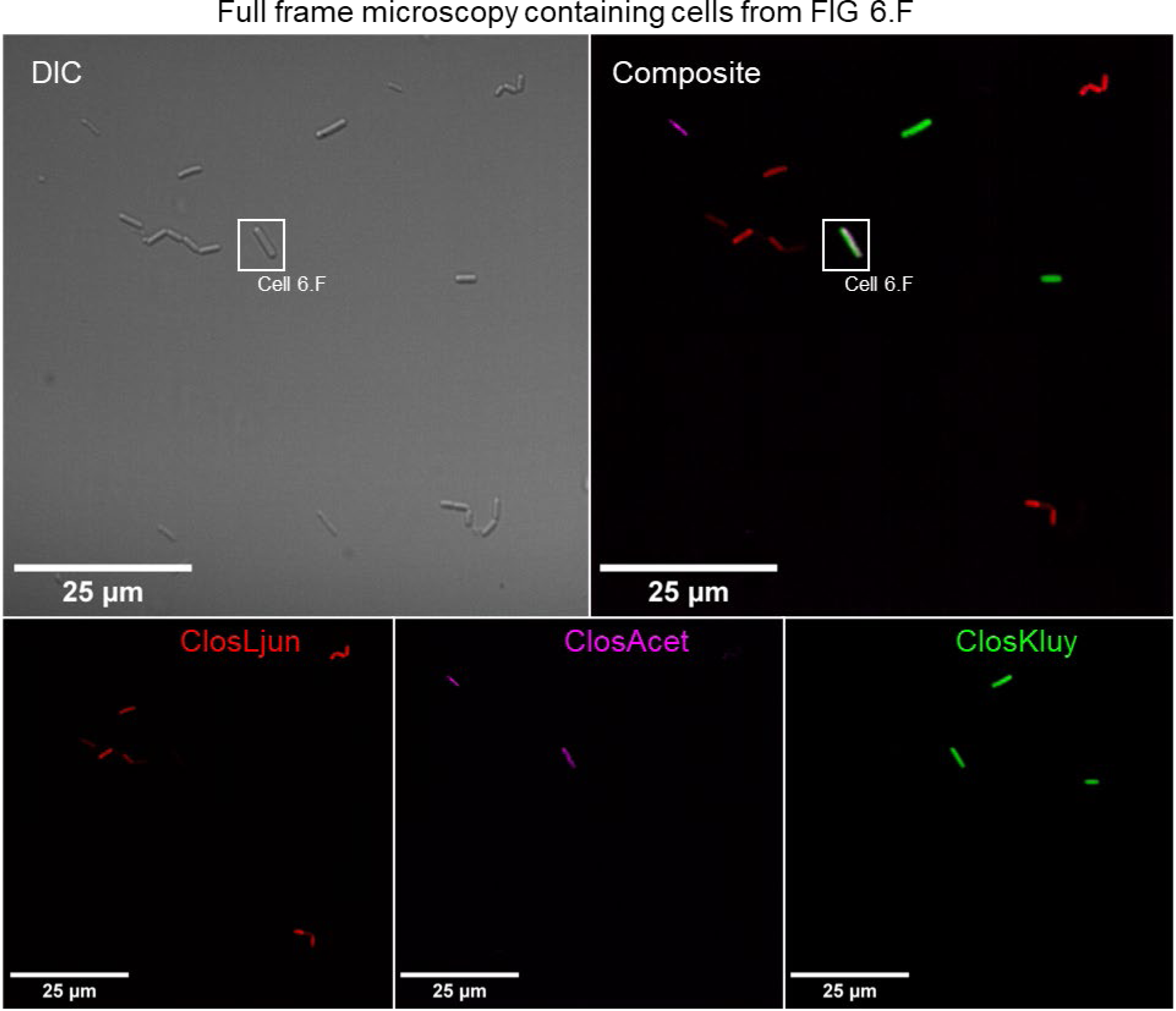
Full Frame images of the cells depicted in Fig. 6F. In the Top row, the DIC and the composite fluorescent image are presented, with the individual fluorescent channels comprising the bottom row.

**Fig S11.**
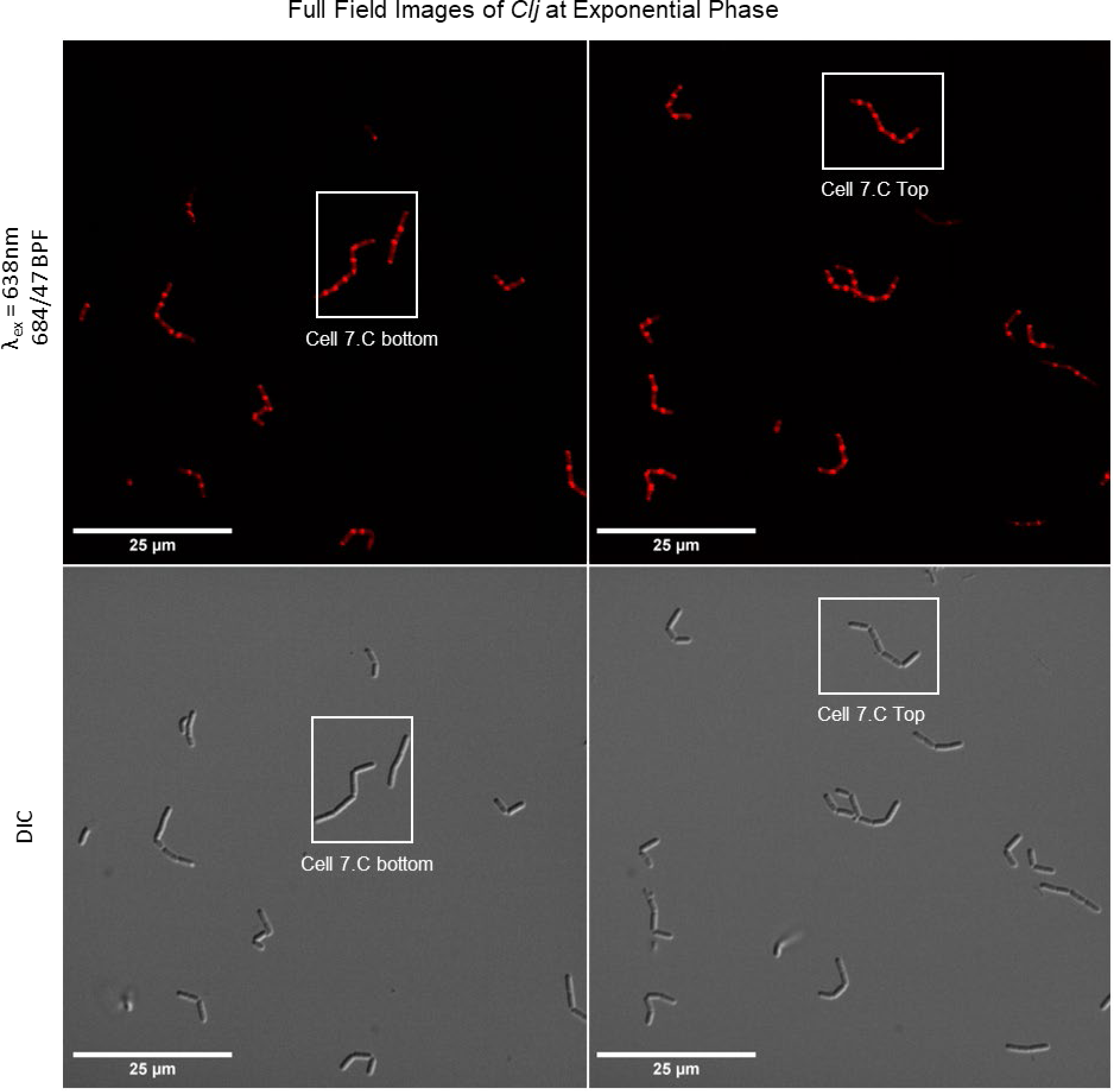
Full frame microscopy images of the exponential phase cells from Fig. 7C. Fluorescent above and DIC below.

**Fig S12.**
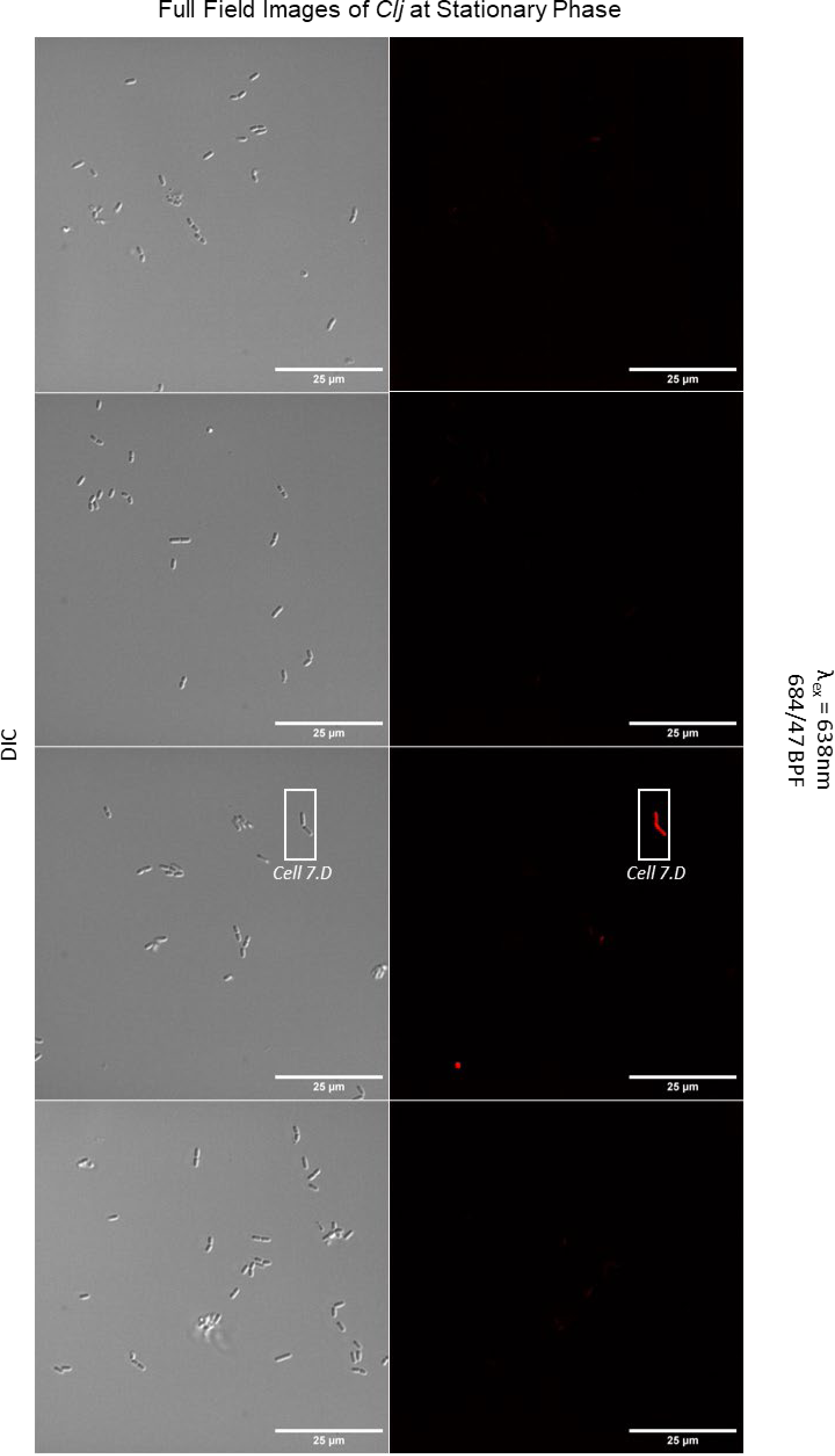
Full frame microscopy images of the stationary phase cells from Fig. 7D. Fluorescent right and DIC left.

**Figure S13.**
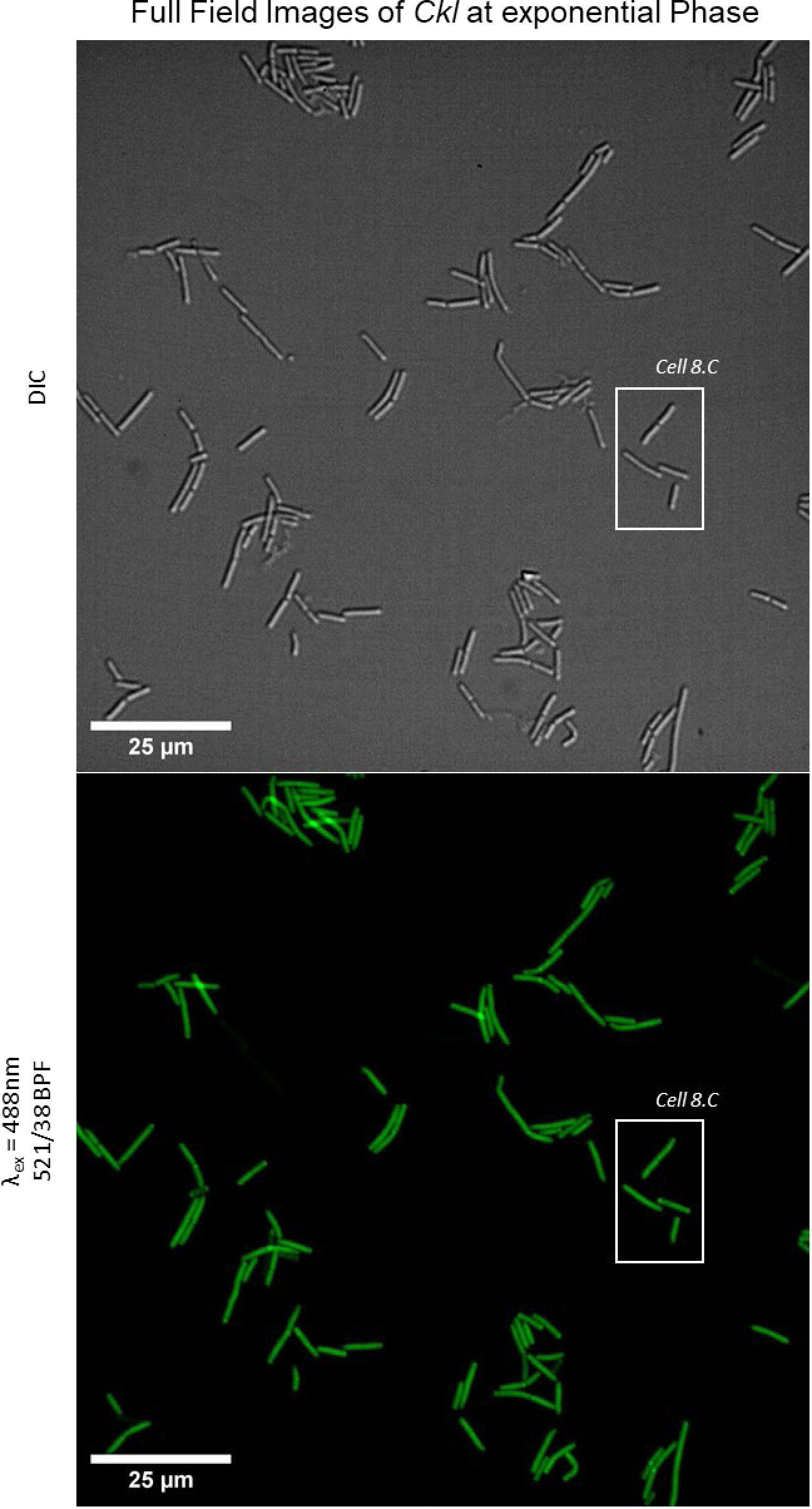
Full frame microscopy images of the exponential phase cells from Fig. 8C. Fluorescent below and DIC above.

**Figure S14.**
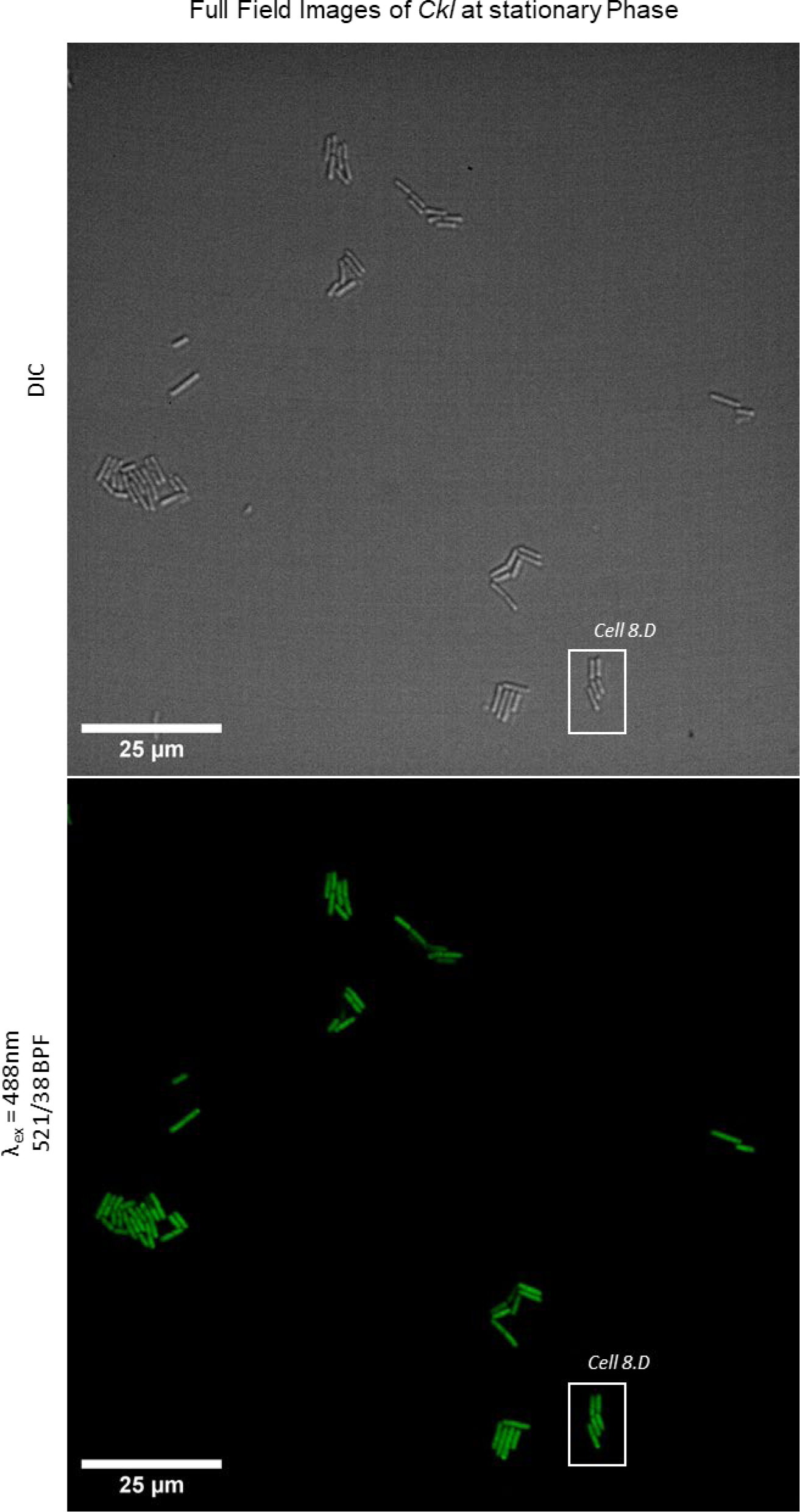
Full frame microscopy images of the stationary phase cells from Fig. 8D. Fluorescent below and DIC above.

**Fig S15.**
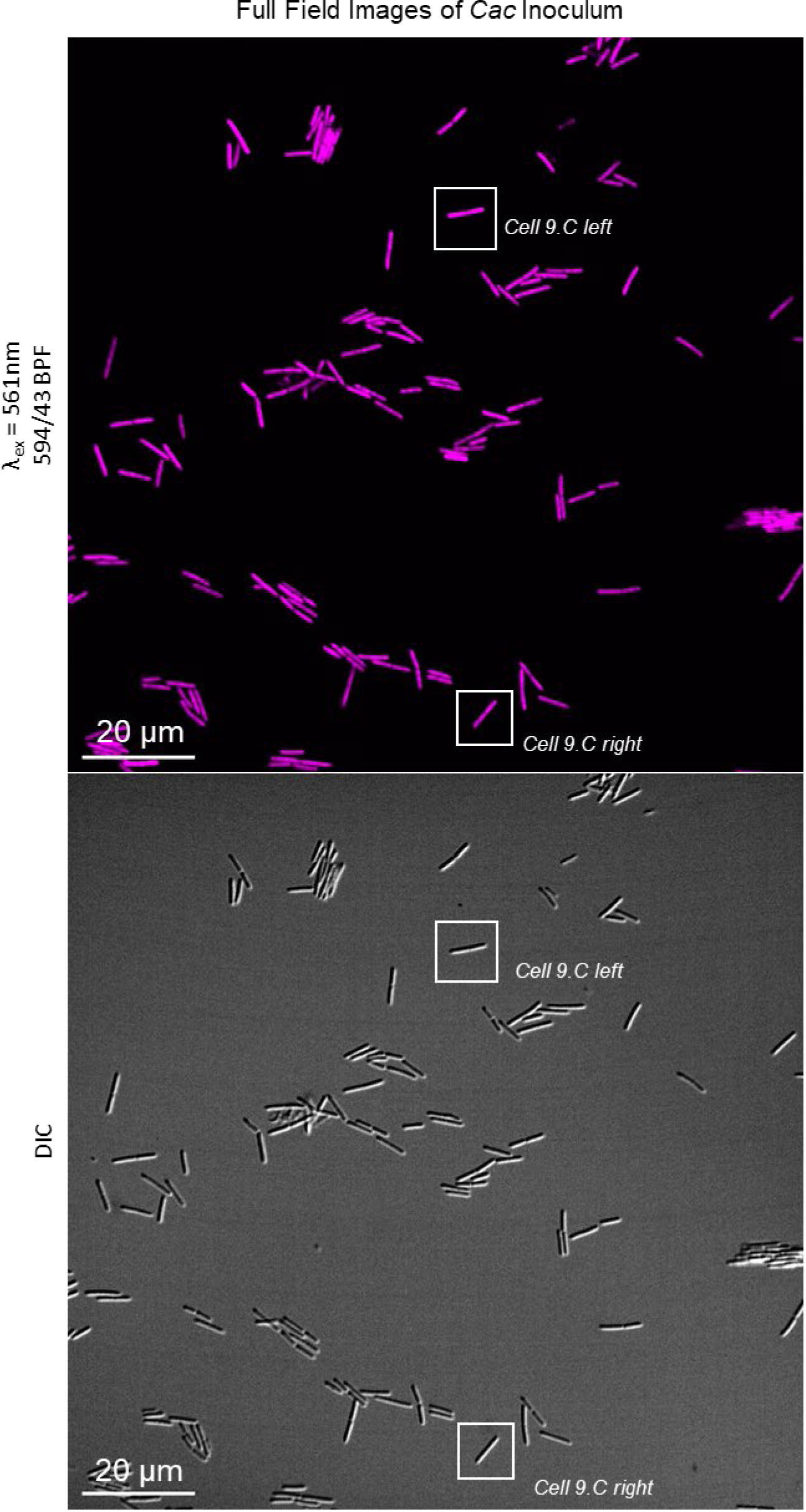
Full frame microscopy images of the inoculum from Fig. 9C. Fluorescent above and DIC below.

**Fig S16.**
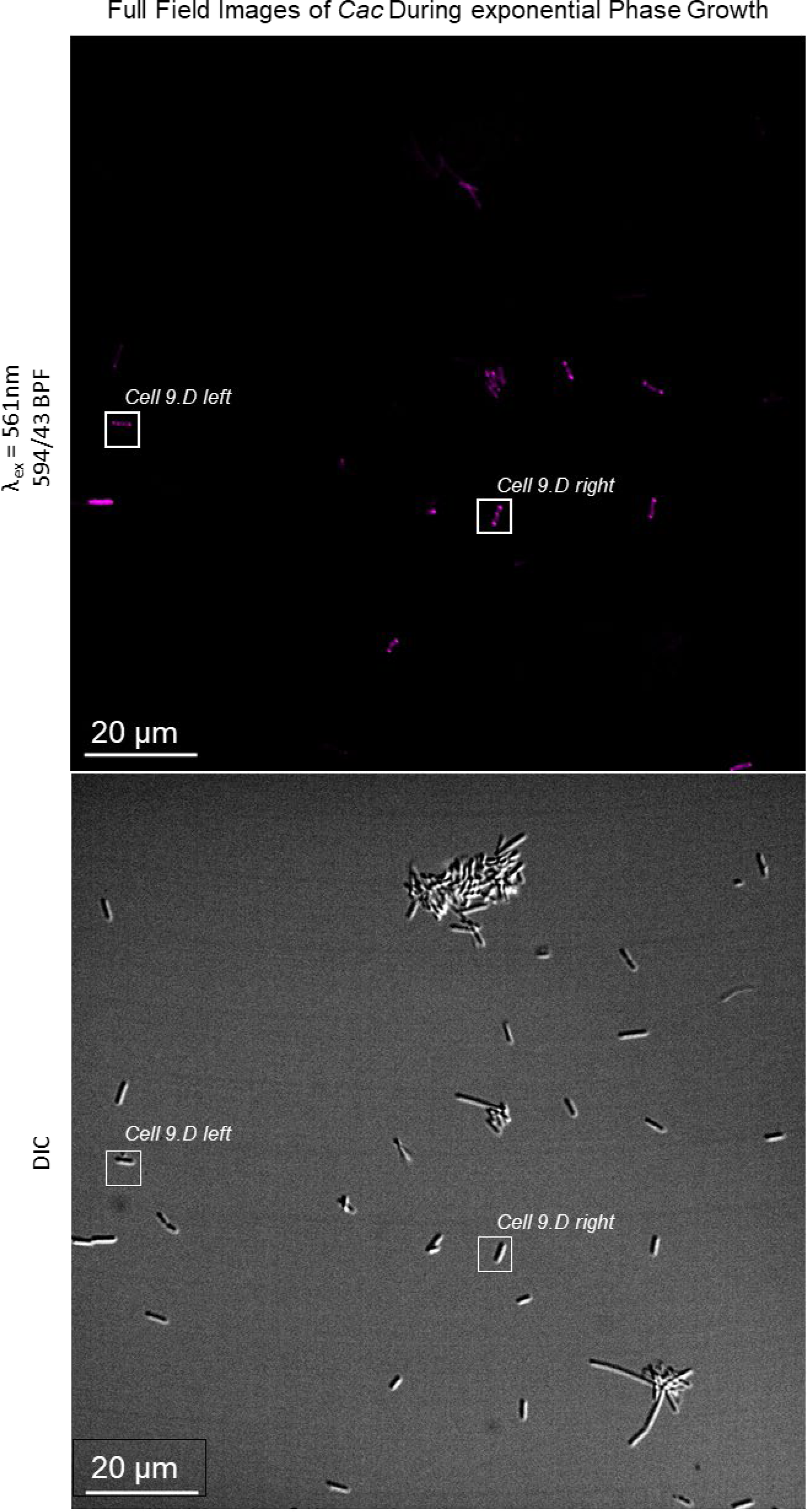
Full frame microscopy images of the exponential phase cells from Fig. 9D. Fluorescent above and DIC below.

**Fig S17.**
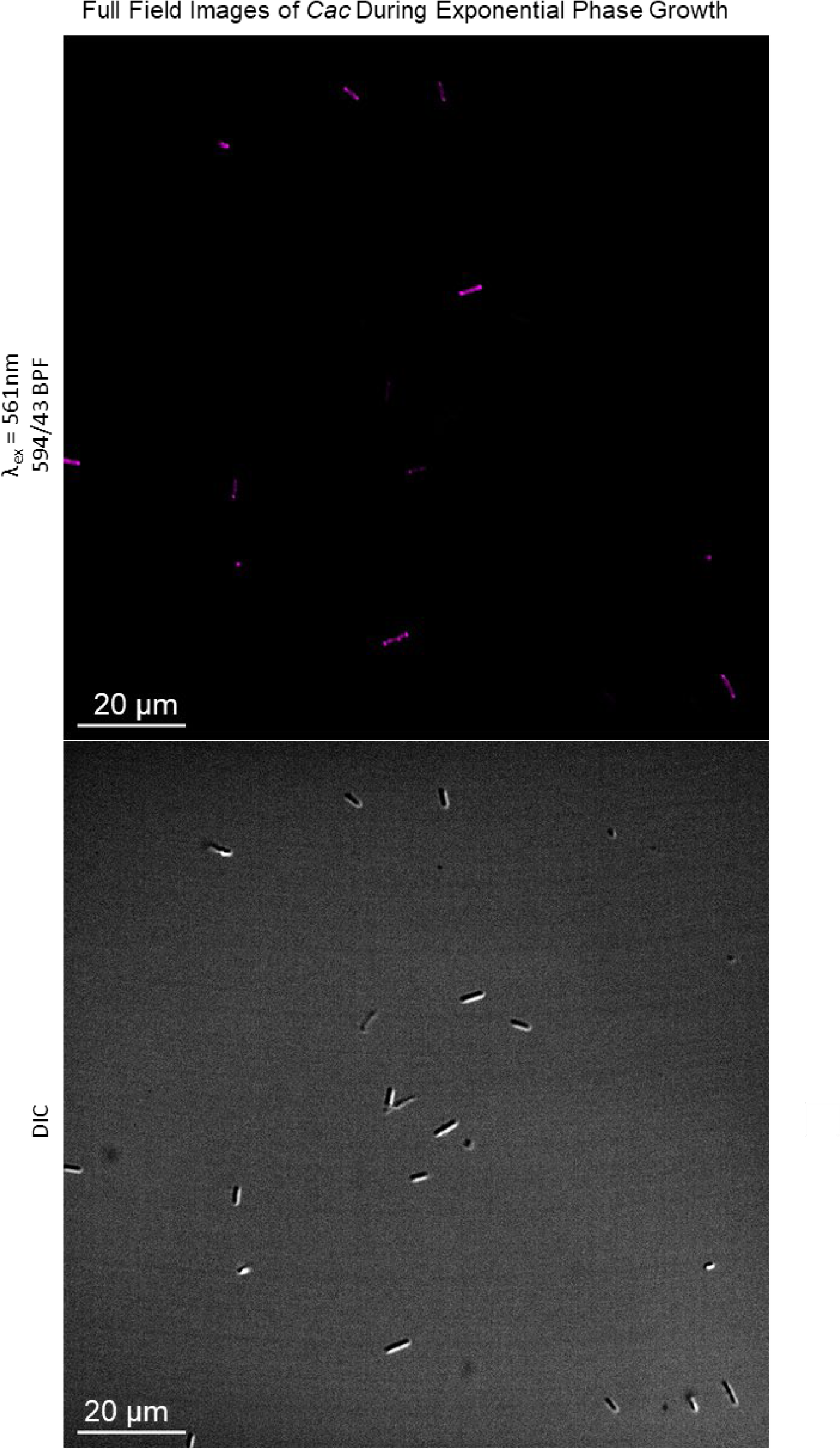
Additional Full frame microscopy images of the exponential phase cells from Fig. 9D. Fluorescent above and DIC below. This second image (in addition to Fig. S16) is included because the cell density is lower.

**Fig S18.**
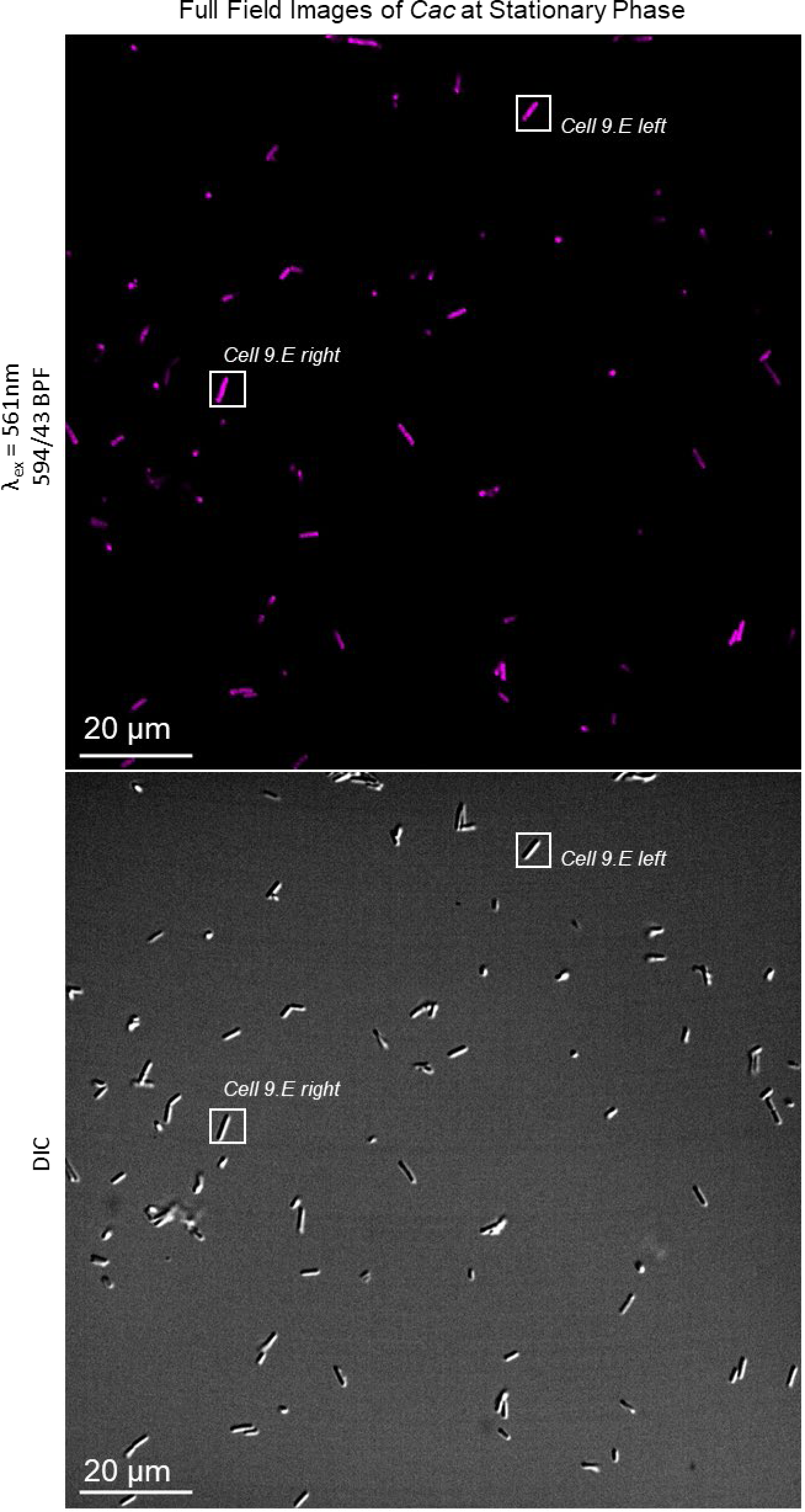
Full frame microscopy images of the stationary phase cells from Fig. 9E. Fluorescent above and DIC below.

